# The order, but not the structure, of cross-domain learning influences memory consolidation

**DOI:** 10.64898/2026.01.13.699035

**Authors:** Ainsley Temudo, Owen Benzley, Bradley R. King, Genevieve Albouy

## Abstract

Everyday activities often require learning sequences that necessitate the involvement of both the declarative and the procedural memory domains. Previous research has shown that a learning structure that is common across tasks from different domains can improve learning and resistance to interference. However, it remains unknown whether such shared learning structure can enhance longer-term memory retention. To address this question, forty-eight healthy adults participated in a pre-registered study in which they learned both an object sequence task (declarative learning) and a motor sequence task (procedural learning) in two separate sessions separated by 4h. Participants were assigned to either an *associated* group, where the two tasks shared a common learning structure - that consisted of a specific mapping between finger movements and object categories across learning sessions - or an *unassociated* group with no such cross-domain shared structure. Memory retention was assessed with a 24h retest session on both tasks. Contrary to our predictions, a shared higher-order structure between tasks from different domains did not enhance memory retention. Exploratory follow-up analyses revealed that the order the tasks were learned (i.e., object or motor first), rather than their structural overlap, influenced performance. Specifically, learning the motor task before the object task *impaired* the consolidation of the object task irrespective of whether the tasks shared a common learning structure or not. This effect was unidirectional as learning the object task before the motor task had no effect on the consolidation of the motor task. Altogether, the current findings suggest that the order of cross-domain learning experiences rather than their structure influences memory consolidation.

## Introduction

Traditionally, memory in humans has been described to be organized in different systems with the declarative system supporting memory of facts and events, and the procedural system supporting the acquisition of movement-based skills [1]. Many everyday behaviors require the integration of information supported by these two distinct systems. For instance, learning to tie shoelaces involves declarative memory to learn the sequence of steps (e.g. make a loop, wrap around, pull through) and procedural memory to develop the motor proficiency required to execute these steps (i.e., the sequence of necessary finger and hand movements). The execution of such behaviors involves sequences of interrelated components — such as individual steps/instructions or actions/movements — that together form a coherent memory and depend on interactions between the memory domains mentioned above.

Although these two memory systems interact continuously in daily life, they have traditionally been studied in isolation [2–5]. This separation was grounded in early neuropsychological findings demonstrating anatomical and functional segregation between domains such that patients with hippocampal lesions displayed deficits in declarative memory but intact procedural learning, while those with striatal dysfunction showed the reverse pattern [1,6,7]. However, these dichotomic views were challenged over the last two decades by both behavioral and neuroimaging studies. The first experimental evidence in support of shared neural resources between memory domains was provided by [8] in a behavioral study showing that memory consolidation in one domain can be disrupted when learning occurs in a different memory domain immediately afterwards [8]; but see [9] for a failed replication). Specifically, participants who learned a motor sequence followed by a declarative wordlist showed impaired motor performance at retest — and the same pattern of interference was observed when the order of learning was reversed — suggesting that declarative and procedural memories share neural resources. This is in line with subsequent neuroimaging studies showing overlapping neural processes between memory domains. For example, and in contrast to the early neuropsychological research cited above that has traditionally associated the hippocampus with the declarative memory domain, several studies have demonstrated that the hippocampus contributes to the learning of motor (procedural) sequences [3,5,10,11]. Altogether, these earlier studies highlight the importance of examining memory processes in a more integrated manner that considers cross-domain overlaps and interactions.

In a series of recent behavioral studies that followed up on the observations reported above, Robertson and collaborators tested whether manipulating cross-domain interactions–with the introduction of higher-order associations between tasks from different domains–could influence the learning process. Higher-order associations between memory domains in this earlier work consisted of creating a common learning structure between a sequence of movements and a sequence of words. Specifically, each finger movement in the motor sequence was linked to a specific word category in the sequence of words, creating a shared structural mapping between the two sequences. Results showed that when the two tasks shared this common temporal structure, learning in one domain facilitated performance in the other domain, suggesting that structural overlap between tasks can promote learning transfer across memory systems [12,13]. In a follow-up study, the authors tested whether such higher-order associations could protect memories against the deleterious effect of interference. Results showed that memory consolidation in one domain was less sensitive to interference provided that participants learned the task from the other domain that shared a common learning structure prior to the interference session [14]. Altogether, these findings indicate that interactions between the declarative and procedural memory systems can modulate the learning process (but see [15] for a failed replication). However, it remains unclear how such cross-domain associations influence longer-term memory retention.

To address this knowledge gap, the present pre-registered study examined whether a shared temporal structure between sequential tasks from different memory domains influences memory retention across an overnight interval. We used an adapted version of the serial reaction time task (SRTT, [11] that allowed us to assess both motor sequence learning (procedural memory) and object sequence learning (declarative memory) in separate versions of the same task. Participants learned both the motor and object sequence tasks in two separate sessions and were divided into two groups according to whether the tasks shared or not a common temporal structure. Based on the previous findings reviewed above, we hypothesized that higher-order associations between the motor and object sequential tasks would result in greater overnight memory consolidation compared to when tasks did not share a common temporal structure.

## Material and Methods

This study was pre-registered in the Open Science Framework (https://osf.io/). Our pre-registration document outlined our hypotheses and intended analysis plan as well as the statistical models used to test our a priori hypotheses (available at https://osf.io/5wmha/). Whenever an analysis presented in the current paper was not pre-registered, it is referred to as *exploratory*.

### Participants

A total of 48 young (aged 18-35) healthy volunteers (30 female, mean age=22 SD=3.7) were recruited for this study. Sample size computation was done with G*Power [16]. Since there was no prior study investigating the effect of higher-order associations between memory domains on memory retention at 24h, the power computation was based on a previous behavioral study investigating the effect of a shared structure on learning transfer across different memory tasks administered within the same day [13]. The effect size was calculated based on the behavioral main effect of interest showing a significant effect of group reflecting learning transfer between tasks from different domains sharing the same structure (different vs. same; Cohen d = 1.23). The estimated sample size was 19 subjects per group (effect size d=1.23, tails=2, alpha=0.05, power=0.95, allocation ratio=1). In our experiment, the condition order (motor or object) and the sequence learned (sequence A or sequence B) were counterbalanced across participants (see procedure below). To balance combinations across participants, the sample size had to be a multiple of 8 (i.e., number of possible condition/sequence combinations), therefore 24 participants were recruited in each group.

Before participating in the study, participants completed an online screening questionnaire to verify their eligibility. The inclusion criteria were: 1) no previous extensive training with a musical instrument requiring dexterous finger movements (e.g., piano, guitar) or as a professional typist, 2) free of medical, neurological, psychological, or psychiatric conditions, including depression and anxiety as assessed by the Beck’s Depression and Anxiety Inventories [17,18], 3) no indications of abnormal sleep, as assessed by the Pittsburgh Sleep Quality Index (PSQI; [19], 4) not considered extreme morning or evening types, as quantified with the Horne & Ostberg chronotype questionnaire [20] and, 5) free of psychoactive and sleep-influencing medications, and 6) normal levels of daytime sleepiness assessed with the Epworth sleepiness scale [21]. Handedness was also assessed using the Edinburgh Handedness Inventory [22]. As per our pre-registration, participants would be excluded if they failed to learn or accurately perform the correct motor and object sequences during the training session on Day 1 (criteria detailed in the behavioral analysis section) and/or if they did not comply with experimental instructions (e.g., maintenance of regular sleep schedule, use of drugs and alcohol; see procedure). However, no participants presented these characteristics and therefore no one was excluded in this study. Participants were recruited through local advertisements posted on the University of Utah campus. All study procedures were approved by the University of Utah institutional review board (IRB_00155080). The study was conducted according to the principles expressed in the Declaration of Helsinki. All participants provided written informed consent at the start of the study and received monetary compensation for their time. Participant demographics as well as information on their sleep and vigilance are presented in the supplemental information (Table S1 and S2).

### Task

All participants performed an adapted version of the explicit Serial Reaction Time Task (SRTT) previously used in our earlier research [11] to examine sequence learning in the two different memory domains (procedural vs. declarative) using the same task probing either motor or object sequence learning. The task was designed such that performance improvement on the motor sequence learning task (as compared to random) reflects sequence-specific skill acquisition and performance improvement on the object sequence learning task (as compared to random) reflects the acquisition of the explicit knowledge of the order of the series of objects (see below for details). The task consisted of a 4-choice reaction time task in which participants were instructed to react to visual cues shown on a screen by pressing a key on a keyboard using their non-dominant hand. A large visual cue shown in the center of the screen corresponded to one of eight different images of objects that belonged to 1 of 4 object categories (animals, vehicles, fruit, and tools). The 8 different objects were constantly presented on the bottom of the screen in 2 rows of 4 objects that mapped to each of the 4 fingers used to perform the task (i.e., no thumbs). Object cues were presented one after the other in the center of the screen and participants were instructed to respond to the cue as fast and as accurately as possible by pressing the corresponding key according to the object/key mapping displayed at the bottom of the screen (Figure 1A). Importantly, the key-object mapping displayed at the bottom of the screen changed after each stream of 8 elements (i.e., every 8 objects or keys) in order to orthogonalize motor and object sequence learning. With this manipulation, three different conditions were developed [11]. In the *random condition*, the series of both responses and objects followed a random order. Performance improvements in this condition were expected to be minimal and represent general visuomotor habituation to the task, given that there was no (object or motor) sequence to learn. In the *object sequence condition*, the stream of objects on the screen followed a repetitive sequential pattern (object sequence, e.g., saw, giraffe, pineapple, car, elephant, axe, plane, banana) while the responses on the keyboard followed a random order. Improvement in performance on this condition (compared to random) was therefore specifically related to the knowledge of the series of objects independent of the series of finger presses. In the *motor sequence condition*, the responses on the keyboard followed a repetitive sequential pattern (motor sequence, e.g., 4,1,3,2,1,4,2,3, whereby 1 and 4 represent the little and index finger, respectively) while the stream of objects on the screen followed a random order. In this condition, performance improvement (as compared to random) therefore specifically reflected the learning of the sequence of movements independent of the series of objects. Accordingly, contrasting performance between the random and sequential SRTT allowed for the extraction of sequence-specific learning in both memory domains (i.e., beyond general visuomotor learning). For all task variants, each block included 40 key presses (5 repetitions of the 8-element sequence) separated by 10-second rest periods. The response stimulus interval (RSI) was 1s where the central object cue was replaced with a green cross. Rest periods were indicated by a red fixation cross in the center of the screen (replacing the central object cue) and red frames replaced the locations of the 8 objects that displayed the key-object mapping at the bottom of the screen. Participants were made aware that the order of objects/movements would follow a repeating sequential pattern but were not told the precise sequence or the number of elements.

**Figure 1.**
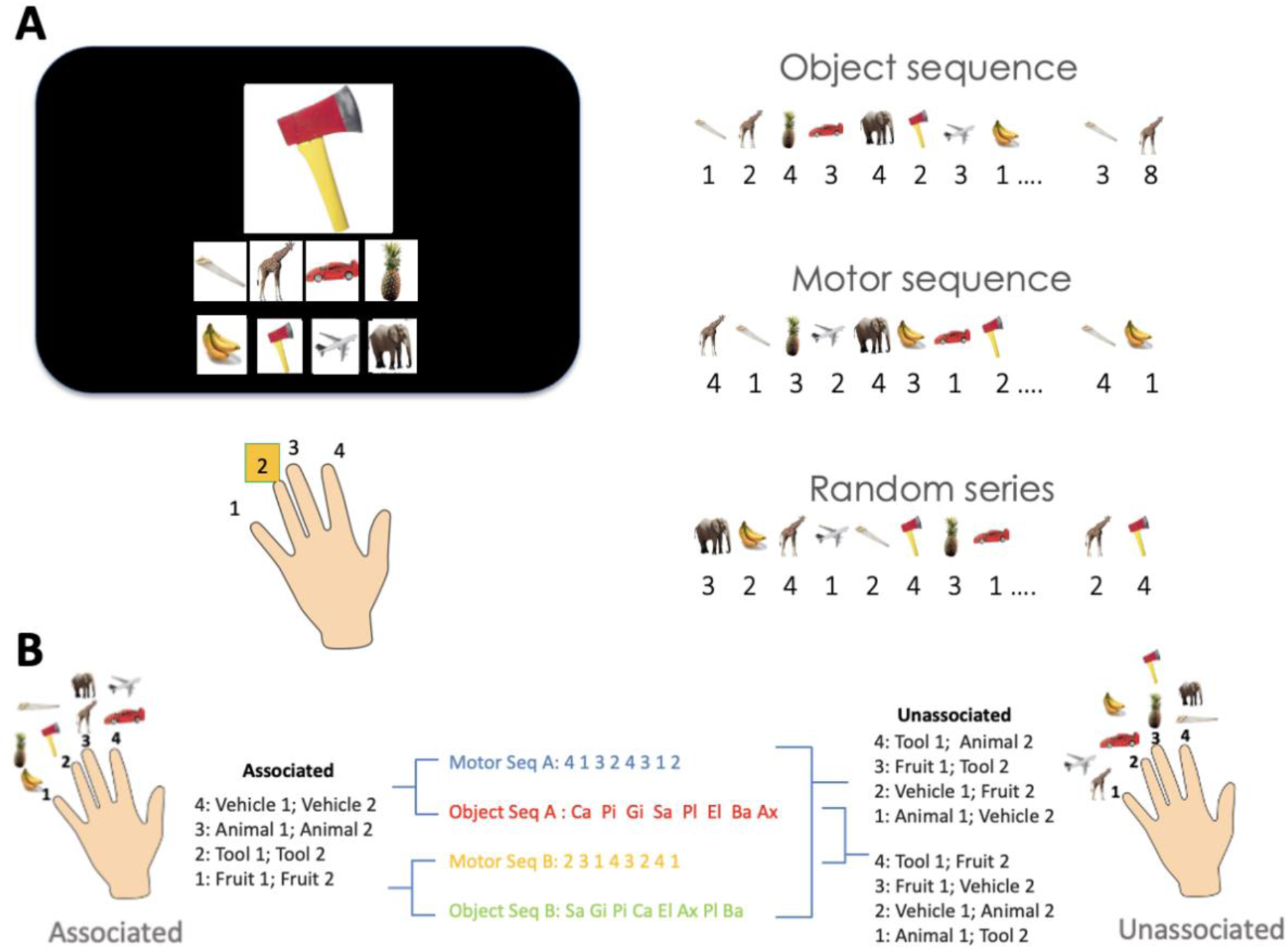
Task. ***A**.* Serial reaction time task (SRTT). Left panel. A stimulus appears in the center of the screen and participants are instructed to respond as fast and as accurately as possible according to the key/object mapping displayed at the bottom of the screen. Note that the key-object mapping changes after each stream of 8 elements (i.e., every 8 objects / 8 key presses) to orthogonalize motor and object sequence learning within each learning session (see right panel). Right panel. Three different task conditions were designed: object sequence with random key presses, motor sequence with random object presentation and a random series with random key presses and random object presentation. Numbers represent fingers (see image of the hand as part of left panel), whereby 1 and 4 correspond to little finger and index finger of the participants’ non-dominant hand respectively. ***B***. Finger-object associations across learning sessions for the associated (left panel) and unassociated (right panel) groups. The middle panel shows the two motor and two object sequences (seq) and the different sequence combinations for each group (associated and unassociated). *Ca=car, Pi=pineapple, Gi=giraffe, Sa=saw, Pl=plane, El=elephant, Ba=banana, Ax=Axe.* All object images were derived from an open access online database at http://olivalab.mit.edu/MM/uniqueObjects.html [24]. The picture of the hands is taken from an open access online sharing service at https://www.clker.com.

Noteworthy, as declarative memory is also defined as the conscious recollection of previous events, we implemented an object generation/recall test aiming at reconstructing the temporal order of the object items as in earlier research [5]. For completeness, a motor generation task was also included to test for any explicit knowledge of the learned sequences of movements as in earlier work [23]. For these generation tasks, participants were instructed to verbally list the objects memorized during the object session and to generate on the keyboard the sequences of movements learned during the motor session. In both generation tasks, participants were asked to generate the respective sequences 3 times in a row, at their own pace, while prioritizing accuracy over speed. Results of the generation tasks are presented in the supplemental text (Section 6). Briefly, generation data indicated that participants presented explicit knowledge of both the sequences of objects and movements.

Importantly, this task paradigm allowed us to examine the effect of higher-order associations between memory domains. Specifically, the task was designed such that there were - or not - associations between fingers and object categories across the motor and object sequence learning tasks (Figure 1B; left and right panels). In the **associated** version of the task, each finger was associated to a particular object category (e.g., key 1: fruit; key 2: tool, key 3: animal and key 4: vehicle) across the motor and object learning sessions on Day 1, while no such finger-object category association was present in the **unassociated** version of the task. Note that within each learning session on Day 1, object and motor items were orthogonalized as described above; therefore, associations between memory domains only occurred across learning sessions on Day 1 and not within each learning session. We designed two different motor sequences (motor sequence A: 4,1,3,2,4,3,1,2 and motor sequence B: 2,3,1,4,3,2,4,1) and two associated object sequences (object sequence A: car, pineapple, giraffe, saw plane, elephant, banana axe and object sequence B: saw, giraffe, pineapple, car, elephant, axe, plane, banana) according to the finger-object category association described above (Figure 1B; middle panel). Motor and object sequences were paired across learning sessions on Day 1 such that specific pairs were “associated” according to the finger-object category association (i.e., motor sequence A associated to object sequence A and motor sequence B associated to object sequence B, Figure 1B left panel) while other pairs were “unassociated” (i.e., motor sequence A unassociated to object sequence B and motor sequence B unassociated to object sequence A, Figure 1B right panel) as they did not follow the finger-object category association mentioned above. Pair combinations were balanced within each group as well as across groups.

### Procedure

Participants completed 3 experimental sessions spread over 2 consecutive days (Figure 2). They were instructed to follow a constant sleep schedule (according to their own schedule ±1h; latest bedtime: 1am; minimum 7h of sleep; no naps) starting 3 days before the first session and between the two experimental days. Compliance to the sleep schedule was monitored using sleep diaries and wrist actigraphy (ActiGraph wGT3X-BT, Pensacola, FL).

**Figure 2.**
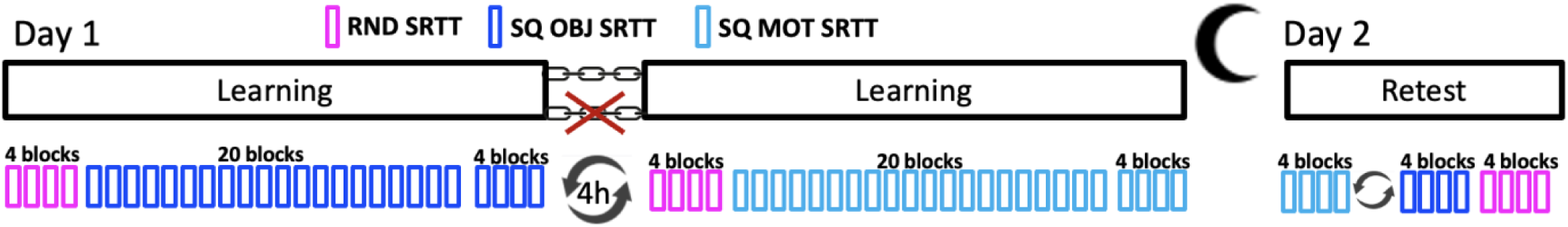
Experimental design. On Day 1, after baseline performance assessment with a random (RND) SRTT, participants learned a sequence of objects (SQ OBJ SRTT) and a sequence of movements (SQ MOT SRTT), in two separate sessions 4h apart (order counterbalanced across participants). On Day 2, a retest session was performed to assess memory retention on each sequence condition (order counterbalanced across participants) followed by a final test on the random condition. Both groups followed the same experimental design; however, the object and motor sequence learning tasks were linked by a higher-order association only in the associated group (Asso; *n* = 24; depicted by a chain link), and not in the unassociated group (Unasso; *n* = 24; depicted by a crossed-out chain link).

On experimental Day 1, participants were invited for 2 experimental sessions separated by 4h. Experimental session 1 was performed between 9am-12pm and started with completing a brief questionnaire to collect information regarding participants’ sleep patterns in the previous 24h (St Mary’s Hospital Sleep Questionnaire [SMS]; [25], their subjective feelings of alertness (Stanford Sleepiness Scale [SSS]; [26], and any consumption of alcohol and/or recreational drugs in the last 24h. Next, participants performed a psychomotor vigilance test (PVT; [27] to obtain an objective measure of alertness. For the PVT, participants were asked to fixate on a fixation cross in the center of the screen and to press the space bar as fast as possible when the fixation cross disappeared. A timer appeared on the screen displaying the participants’ reaction time in milliseconds. Participants completed 50 trials of the PVT task. Participants were then invited to practice the SRT task. First, participants completed 4 blocks of the random condition to get familiar with the task apparatus and measure baseline performance. They were then trained on 20 blocks of either the object sequence (A or B) or the motor sequence (A or B). After a short break offered to minimize the confounding effect of fatigue on end-training performance [28], performance was tested with 4 additional blocks of practice. After completing experimental session 1, participants were given actigraph monitors to be worn until the end of the experiment. For the 4h between experimental sessions 1 and 2 on Day 1, participants were instructed to not take a nap or practice the task and to avoid intense exercise and consumption of substances with alcohol or caffeine.

Experimental session 2 took place at least 4h after session 1 (i.e., between 1-4pm) in order to reduce interference between the 2 learning episodes [8,13]. This session followed the same procedure as session 1, except that participants were trained on the other sequential condition (object or motor). The order of task conditions (object vs. motor) and sequence types (A vs. B) was counterbalanced across participants. The pair of sequences attributed to each individual depended on their group assignment (motor sequence A - object sequence A and motor sequence B - object sequence B for participants in the **associated group** while other pairs such as motor sequence A - object sequence B and motor sequence B - object sequence A for participants in the **unassociated group**). After completing the experimental session 2 on Day 1, participants were instructed to have a regular night of sleep according to the schedule followed prior to the experiment, to continue wearing the actigraph, to not practice the task and to avoid consumption of substances with alcohol or caffeine.

Participants returned on Day 2, approximately 24h after session 1, for the third and final experimental session. Participants were asked to complete the same questionnaires that were completed at the start of experimental session 1 on Day 1. Next, they performed the PVT followed by 4 blocks of each of the 2 sequential conditions in a counterbalanced order. A final 4 blocks of the random condition were performed to assess sequence-specific memory retention. Experimental session 3 ended with the generation tasks described above to test whether participants presented explicit awareness of the sequences.

### Behavioral data analysis

#### Preprocessing & Quantification

During performance of the SRT tasks, the keys pressed, and timing of the key presses were recorded, and performance was quantified using **speed** and **accuracy** measures. Specifically, performance speed was defined as the time between the visual cue onset and the corresponding response (i.e., response time). For accuracy, a trial was classified as “correct” if the key pressed by the participants matched the object cue (i.e., a correct response), and as “incorrect” if it did not (i.e., an incorrect response). The averaged response times for correct key presses and the percentage of correct key presses were computed for each block of practice for each version of the task. Consistent with our pre-registration, individual trials (i.e., responses) were excluded from the analyses if the measured response time was greater than 3 standard deviations above or below the participant’s mean response time for a given block (1.63 % of trials excluded across participants). Results of the analysis performed on the speed measure are presented in the main text while those related to the accuracy measure are presented in the supplementary text.

To examine the effect of cross-domain higher-order associations on consolidation processes, we computed the **offline changes in performance** between Day 1 and Day 2. To do so, the four test blocks at the end of training session 1 and session 2 (Day 1) were averaged separately to calculate **end-of-training performance** on both task conditions and the four test blocks for each task condition in session 3 (Day 2) were also averaged separately for each condition to calculate **retest performance**. Offline changes in performance reflecting the consolidation process were computed as the difference between (averaged) end-of-training and retest performance for each task condition. For the generation tasks, the percentage of items (keys or object) placed in their correct ordinal position as well as the percentage of correct transitions (i.e., two correct consecutive items) were computed for each condition as the average across the three attempts (results of the generation task analyses are reported in Section 6 of the supplemental information).

#### Confirmatory analyses

Our primary confirmatory analysis was to examine whether a common learning structure between tasks from different domains facilitates the memory consolidation process. As pre-registered, we examined offline changes in performance speed and accuracy using repeated-measures ANOVAs with the within-subject factor *session* (end of training vs. retest) and between-subject factor *group* (associated vs. unassociated). Results on accuracy are presented in Section 3 of the supplemental information and did not show any significant group effect.

#### Negative control analyses

##### Initial learning (Day 1)

To confirm that any observed group differences in the results of our confirmatory analysis could be attributed to our experimental manipulation, a series of control analyses were conducted on the training data acquired on Day 1. First, baseline motor execution was compared between associated and unassociated groups using separate repeated-measures ANOVAs on performance (speed and accuracy) of the random serial reaction time task. In one analysis, *block* (1–4) and *session* (1 vs. 2, irrespective of task condition) were entered as within-subject factors and *group* as a between-subject factor. In a separate analysis, *task condition* (motor vs. object, irrespective of session) replaced session as a within-subject factor. Last, an ANOVA using both *block* and *session* as within subject factors as well as *task order* (motor first vs. object first) and group as between-subject factors assessed the effect of *task order* on random performance. These results are reported in Section 1 of the supplemental information and collectively show that baseline performance on a random SRTT was similar between groups and task conditions but tended to be faster and less accurate in session 2 as compared to session 1. They also show an effect of task order on random performance (see discussion).

Second, performance (speed and accuracy) on the sequential SRTT on Day 1 was analyzed using repeated-measures ANOVAs conducted separately for training and test data. Four models were tested: (1) an ANOVA using both *block* (20 for training and 4 for test) and *session* (1 vs. 2) as within-subject factors and *group* (associated vs. unassociated) as between-subject factor assessed potential baseline differences between *sessions* independent of *task condition* (motor vs. object), (2) an ANOVA with both *block* and *task condition* as within subject factors as well as group as between-subject factor assessed baseline differences in *task conditions* independent of *session*, (3) an ANOVA using both *block* and *session* as within subject factors as well as *task order* (motor first vs. object first) and group as between-subject factors assessed the effect of *task order* on performance, (4) an ANOVA conducted separately within each *task condition* using *block* as a within subject factor and *sequence type* (Sequence A vs. Sequence B) and *group* as between subject factors, tested for potential differences in baseline performance related to the type of sequence. The results of the last model are presented in Tables S3 and S4 of the supplementary information and indicate no baseline differences in performance between Sequences A and B.

The results of these negative control analyses described above but conducted on the accuracy measure are presented in Section 4 of the supplementary information. Briefly, results indicate that participants were more accurate on the motor task compared to object task and during the first session as compared to the second session, but that accuracy remained similar between groups.

##### Retest (Day 2)

As per the pre-registration, to determine whether any group differences observed on task performance on Day 2 were specific to sequence learning rather than general performance enhancements, we compared performance between the sequence and random task conditions from Day 2 using repeated-measures ANOVAs with *block* (1-4) and *task condition* (motor, object, random) as within-subject factors as well as *group* as a between-subject factor. Results are reported in Section 2 of the supplemental information and indicate that memory retention was sequence-specific and that this effect did not differ between the associated and unassociated groups.

##### Generation

Sequence awareness was assessed from the generation task by comparing the proportion of items in the correct ordinal position and the proportion of correct transitions between task conditions and groups, using repeated-measures ANOVAs with within-subject factor *task condition* (object vs. motor) and between-subject factor *group* (associated vs. unassociated). Results are reported in Section 6 of the supplemental information and indicate that participants developed explicit knowledge of the sequence of objects and movements in both experimental groups.

#### Exploratory analyses

In contrast to our expectations, some of the negative control analyses described above showed trends for baseline performance differences on Day 1. We therefore performed the exploratory analyses described below to further examine these trends. The results of the exploratory analyses on accuracy are presented in Section 5 of the supplementary information and didn’t show any significant effects.

##### Initial learning (Day 1)

To further characterize the session x task order x group effects, we compared task conditions within each session (i.e., same task order, different conditions) between groups using repeated-measures ANOVA including *block* as within-subject factor as well as *condition* and *group* as between-subject factors. A second exploratory analysis was performed within each task condition (i.e., same task condition, different order) with a repeated-measures ANOVA using *block* as within-subject factor as well as *order* and *group* as between subject factors.

##### Gains in performance

We examined whether gains varied as a function of task condition and the order in which the tasks were learnt on Day 1. To do so, we conducted repeated-measures ANOVAs separately within each task condition with *order* (e.g. object session 1 vs. object session 2) and *group* (associated vs. unassociated) as between subject factors. Additional analyses were performed separately for gains from session 1 and session 2 with *condition* (motor vs. object) and *group* as between subject factors.

## Results

### Effect of higher-order associations on memory consolidation

Our pre-registered confirmatory analyses assessed the effect of higher-order associations on memory consolidation. A repeated-measures ANOVA conducted on performance speed averaged across the 4 blocks of post-training test on Day 1 for each condition and the 4 blocks of retest on Day 2 for each condition using *session* (Day 1 vs. Day 2) as within-subject factor and *group* (associated vs. unassociated) as between-subject factor was performed. Results showed a significant effect of session, whereby performance improved from the end of training to retest (session: *F*(1, 46) = 21.14, *p* < .001, ɳp² = .32). In contrast to our expectations, this session effect did not differ between the 2 groups (session x group: *F*(1, 46) = .07, *p* = .80, ɳp² = .001; group: *F*(1, 46) = .20, *p* = .66, ɳp² = .004), indicating that the presence of a higher-order associations between sequences from the two different domains did not enhance offline memory consolidation (Figure 3B).

**Figure 3.**
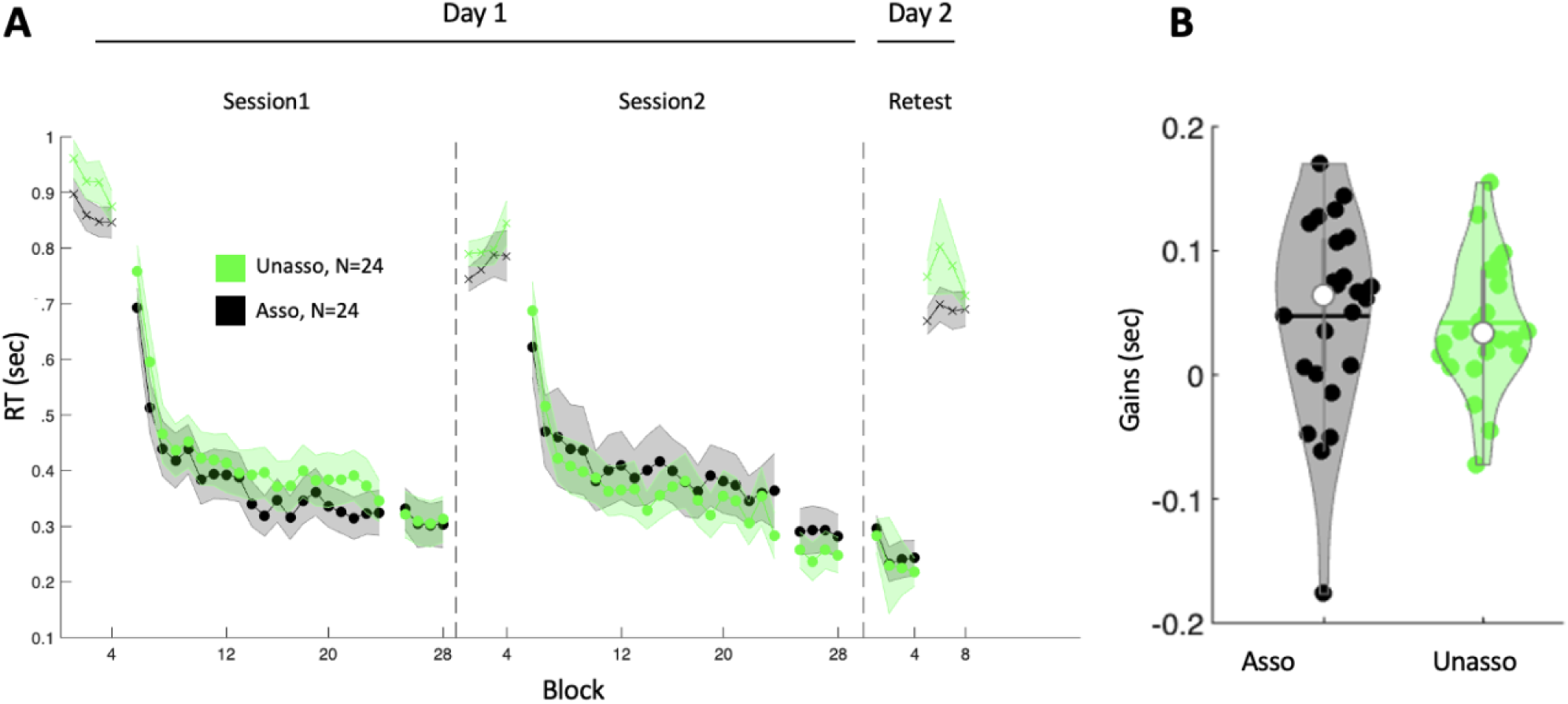
Performance speed results. **A**. Response time (RT, in seconds) across blocks and sessions collapsed across motor and object task conditions for the unassociated (“Unasso”, green, N = 24) and associated (“Asso”, black, N = 24) groups. Crosses and circles indicate blocks of the random and sequence conditions, respectively. Shaded error bars=SEM. **B**. Gains in performance speed (in seconds) calculated as the average response time across the four test blocks on Day 1 minus the average across the four retest blocks on Day 2, for each group. Gains were not significantly different between the associated and unassociated group indicating that higher-order associations between sequences across different memory domains did not enhance offline memory consolidation. Colored circles represent individual data, jittered in arbitrary distances on the x-axis to increase perceptibility. White circles represent medians, and horizontal bars represent means. The shape of the violin depicts the kernel density estimate of the data.

### Negative control analyses

To ensure that the results presented above were not influenced by differences in performance during initial training, we tested for potential group effects during initial task practice on Day 1.

Results of the block x session (session 1 vs. session 2) x group (associated vs. unassociated) repeated-measures ANOVA (Figure 3A) showed a significant block effect, whereby speed improved across training blocks (block: *F*(19, 874) = 48.84, *p* < .001, ɳp² = .52). As expected, performance did not differ between groups (group: *F*(1, 46) = .03, *p* = .88, ɳp² = .001; block x group: *F*(19, 874) = 1.18, *p* = .27, ɳp² = .03) or between sessions (session: *F*(1, 46) = .07, *p* = .79, ɳp² = .002; block x session: *F*(19, 874) = 1.51, *p* = .07, ɳp² = .03) and there was no interaction between these factors (session x group: *F*(1, 46) = .77, *p* = .38, ɳp² = .02; block x session x group: *F*(19, 874) = .64, *p* = .87, ɳp² = .01). Performance speed stabilized at the test (block: *F*(3, 138) = 1.29, *p* = .28, ɳp² = .03) and remained similar between groups (group: *F*(1, 46) = .21, *p* = .65, ɳp² = .005; block x group: *F*(3, 138) = .25, *p* = .86, ɳp² = .005) and sessions (session: *F*(1, 46) = 1.61, *p* = .21, ɳp² = .03; block x session: *F*(3, 138) = .79, *p* = .50, ɳp² = .02; session x group: *F*(1, 46) = .40, *p* = .53, ɳp² = .009; block x session x group: *F*(3, 138) = .46, *p* = .71, ɳp² = .01). These results indicate that learning was similar during both experimental sessions on Day 1 between groups.

The block x condition (motor vs. object, irrespective of session) x group (associated vs. unassociated) repeated-measures ANOVA showed a significant condition effect, whereby the motor task was performed faster than the object task (training; condition: *F*(1, 46) = 29.12, *p* < .001, ɳp² = .39; block x condition: *F*(19, 874) = 2.63, *p* < .001, ɳp² = .05; test; condition: *F*(1, 46) = 30.18, *p* < .001, ɳp² = .40; block x condition: *F*(3, 138) = 1.78, *p* = .15, ɳp² = .04). However, this effect did not interact with the factor group (training: condition x group: *F*(1, 46) = 1.03, *p* = .31, ɳp² = .02; block x condition x group: *F*(19, 874) = .98 *p* = .48, ɳp² = .02; test; condition x group: *F*(1, 46) = .39, *p* = .54, ɳp² = .008; block x condition x group: *F*(19, 874) = .78 *p* = .51, ɳp² = .02). These results indicate that participants perform the motor sequence task faster than the object sequence task which is consistent with our earlier work [11]. Importantly, this effect did not interact with the factor group.

We then tested whether the order of conditions across sessions influenced performance differently between groups. To do so, a block x session x task order (motor first vs. object first) x group repeated-measures ANOVA was performed (Figure 4A). Results showed that task order did not differently influence performance between groups (training: task order x group: *F*(1, 44) = 2.14, *p* = .15, ɳp² = .05; session x task order x group: *F*(1, 44) = 1.02, *p* = .32, ɳp² = .02; block x session x task order x group: *F*(19, 836) = .99, *p* = .47, ɳp² = .02; test: task order x group: *F*(1, 44) = 1.31, *p* = .26, ɳp² = .03; session x task order x group: *F*(1, 44) = .40, *p* = .53, ɳp² = .009; block x session x task order x group: *F*(3, 132) = .77, *p* = .51, ɳp² = .02). Even though the order of the different conditions did not affect performance differently between groups (p = .15, see Figure 4A), we ran exploratory analyses to further examine condition effects within (same order) and between (different order) sessions and the impact of task order on offline changes in performance.

**Figure 4.**
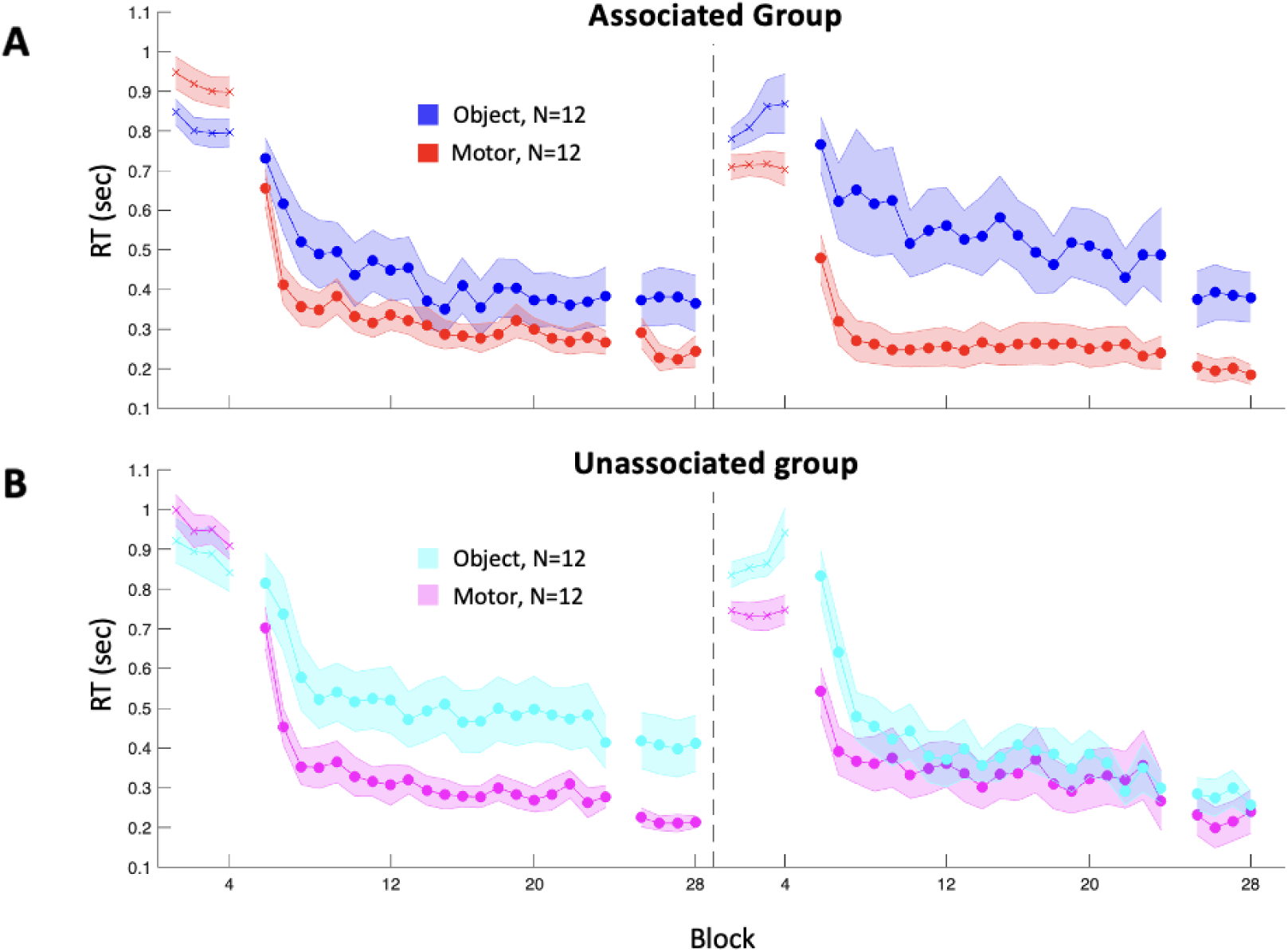
Results of the exploratory analyses on performance speed. **A**. Response time (RT, in seconds) across blocks in session 1 and 2 for the object condition (dark blue, N = 12) and motor condition (red, N = 12) in the associated group. **B**. Response time across blocks in session 1 and 2 for the object (cyan, N = 12) and motor condition (pink, N = 12) in the unassociated group. Crosses indicate blocks of the random condition; circles indicate blocks of the sequence condition. Shaded error bars=SEM.

### Exploratory Analyses

#### Task condition effects within and between sessions (Day 1)

To better understand the potential task order by group effect mentioned above, we compared performance between conditions and groups within each session (different conditions, same order). The block x condition x group RM ANOVA conducted on ***session 1*** (Figure 4A) data confirmed a significant condition effect, whereby the motor task was performed faster than the object task (training; condition: *F*(1, 44) = 7.95, *p* = .007, ɳ_p_^2^= .15; block x condition: *F*(19, 836) = 1.30, *p* = .17, ɳ_p_^2^= .03; test; condition: *F*(1, 44) = 9.75, *p* = .003, ɳ_p_^2^= .18; block x condition: *F*(3, 132) = 1.14, *p* = .34, ɳ_p_^2^= .03). However, this effect did not differ between the associated and non-associated groups (training; group: *F*(1, 44) = .60, *p* = .44, ɳ_p_^2^= .01; condition x group: *F*(1, 44) = .61, *p* = .44, ɳ_p_^2^= .01; block x group: *F*(19, 836) = .78, *p* = .74, ɳ_p_^2^= .02; block x condition x group: *F*(19, 836) = .68, *p* = .84, ɳp^2^= .02; test; group: *F*(1, 44) = .00, *p* = .98, ɳp^2^= .00; condition x group: *F*(1, 44) = .40, *p* = .53, ɳ_p_^2^= .009; block x group: (3, 132) = .27, *p* = .85, ɳ_p_^2^= .006; block x condition x group: *F*(3, 132) = 1.28, *p* = .28, ɳ_p_^2^= .03).

The block x condition x group RM ANOVA conducted on ***session 2*** revealed a similar condition effect (Figure 4A; training; condition: *F*(1, 44) = 7.83, *p* = .008, ɳ_p_^2^= .15; block x condition: *F*(19, 836) = 2.23, *p* = .002, ɳ_p_^2^= .05; test; condition: *F*(1, 44) = 6.71, *p* = .01, ɳ_p_^2^= .13; block x condition: *F*(3, 132) = .79, *p* = .50, ɳ_p_^2^= .02), but here the condition x group interaction showed a trend that did not reach significance (train: *F*(1, 44) = 2.75, *p* = .10, ɳ_p_^2^= .06). This trend was driven by a large condition effect during session 2 in the ***associated group*** (Figure 4B) where performance for the motor condition was faster than for the object condition (training; condition: *F*(1, 22) = 7.54, *p* = .01, ɳ_p_^2^= .26; block x condition: *F*(19, 418) = 1.41, *p* = .12, ɳ_p_^2^= .06; test; condition: *F*(1, 22) = 7.10, *p* = .01, ɳ_p_^2^= .24; block x condition: *F*(3, 66) = .30, *p* = .82, ɳ_p_^2^= .01). This condition effect was not observed in the ***unassociated group*** (Figure 4C; training; condition: *F*(1, 22) = .95, *p* = .34, ɳ_p_^2^= .04; block x condition: *F*(19, 418) = 1.96, *p* = .009, ɳ_p_^2^= .08; test; condition: *F*(1, 22) = .83, *p* = .37, ɳ_p_^2^= .04; block x condition: *F*(3, 66) = 1.38, *p* = .26, ɳ_p_^2^= .06). There was no main effect of group and all other interactions were not significant (training; group: *F*(1, 44) = .18, *p* = .68, ɳ_p_^2^= .004; block x group: *F*(19, 836) = 1.05, *p* = .40, ɳ_p_^2^= .02; block x condition x group: *F*(19, 836) = 1.24, *p* = .22, ɳ_p_^2^= .03; test; group: *F*(1, 44) = .72, *p* = .40, ɳ_p_^2^= .02; condition x group: *F*(1, 44) = 1.89, *p* = .18, ɳ_p_^2^= .04; block x group: *F*(3, 132) = .42, *p* = .74, ɳ_p_^2^= .01; block x condition x group: *F*(3, 132) = .95, *p* = .33, ɳ_p_^2^= .02). Overall, these findings suggest that higher-order associations may have influenced performance of the task learned second during session 2.

A visual inspection of Figure 4B suggests that the results reported above may be driven by slower performance on the object task when it is learned in session 2 as compared to session 1. To test for this hypothesis, we performed a RM ANOVA using *block* as a within-subject factor as well as *order* (object first vs. object second) and *group* as between subject factors (same condition, different order). Results showed a non-significant group by order interaction (training; group x order: *F*(1, 44) = 2.26, *p* = .14, ɳ_p_^2^= .05). There was a trend for slower performance on the object task in session 2 compared to session 1 in the associated group (Figure 4B) and faster performance in session 2 compared to session 1 in the unassociated group (Figure 4C). Note that these effects did not reach significant within each group (associated group, training; order: = *F*(1, 22) = .89, *p* = .36, ɳ_p_^2^= .04; block x order: *F*(19, 418) = 1.08, *p* = .37, ɳ_p_^2^= .05; test; order: = *F*(1, 22) = .007, *p* = .93, ɳ_p_^2^= .00; block x order: *F*(3, 66) = .06, *p* = .98, ɳ_p_^2^= .003; unassociated group, training; order: = *F*(1, 22) = 1.57, *p* = .22, ɳ_p_^2^= .07; block x order: *F*(19, 418) = .73, *p* = .40, ɳ_p_^2^= .03; test; order: = *F*(1, 22) = 2.70, *p* = .12, ɳ_p_^2^= .11; block x order: *F*(3, 66) = .70, *p* = .56, ɳ_p_^2^= .03). No such trends were observed for the motor task (training; order x group: *F*(1, 44) = .86, *p* = .36, ɳ_p_^2^= .02; test; order x group: *F*(1, 44) = .69, *p* = .41, ɳ_p_^2^= .02). Altogether, slower performance in the associated group on the object task in session 2 might reflect a trend for pro-active interference from the motor task (learned first) to the declarative task (learned second). Such interference was not observed when the tasks did not share a common structure or when the object task was learned before the motor task.

#### Gains in performance

As the exploratory analyses presented above suggest that higher-order associations tended to result in interference from the motor to the object task, we examined the effects of these interactions on offline changes in performance.

First, we tested whether a similar order effect was observed on offline changes in performance within each task condition. The 2 (task order) x 2 (group) ANOVA conducted on offline changes in performance speed on the ***object condition*** (Figure 5A) showed a significant order effect (order: F(1, 44) = 5.55, p = .02, ɳp² = .11), whereby gains in performance were lower when the object task was performed in session 2 as compared to session 1. This effect did not differ between groups (group: F(1, 44) = .27, p = .61, ɳp² = .006; order x group; F(1, 44) = .06, p = .81, ɳp² = .001). No such order effect was observed for the ***motor condition*** (Figure 5B; order: F(1, 44) = .32, p = .58, ɳp² = .007; group: F(1, 44) = .05, p = .83, ɳp² = .001; order x group: F(1, 44) = .28, p = .60, ɳp² = .006). These results suggest that learning the motor task prior to the object task might have resulted in a disruption of the consolidation of the object task irrespective of higher-order associations.

**Figure 5.**
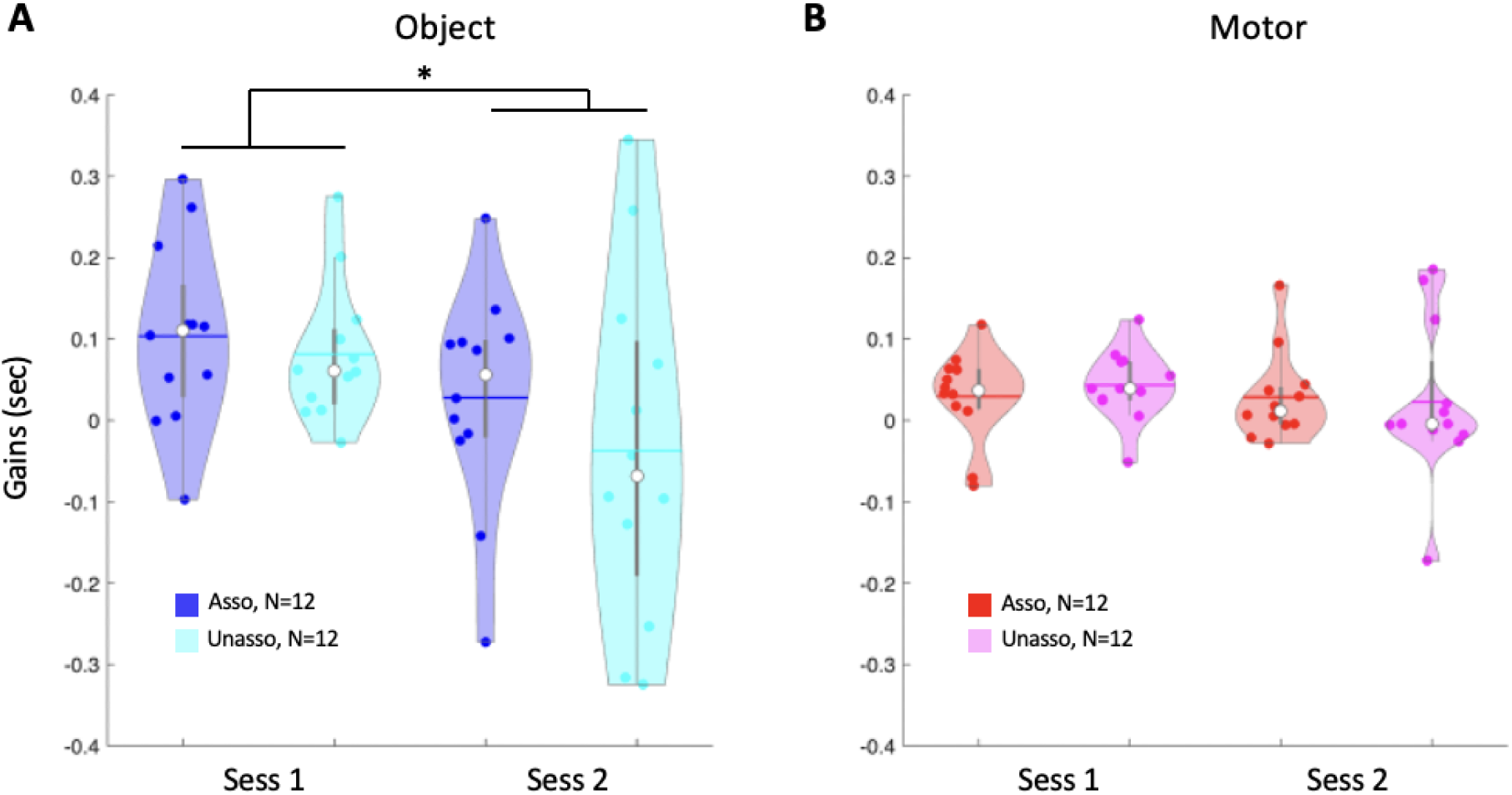
Gains in performance speed. **A**. Gains on the object task in each group depending on whether the object task was learned during session 1 or session 2. **B.** Gains on the motor task in each group depending on whether the motor task was learned during session 1 or session 2. In both panels, gains were calculated as the average response time across the four test blocks on Day 1 minus the average across the four retest blocks on Day 2. Gains in performance speed were lower when the object task was performed in session 2 as compared to session 1 (main effect of order: F(1, 44) = 5.55, p = .02, ɳp² = .11), but not different between groups. No differences in gains on the motor task were observed. Colored circles represent individual data, jittered in arbitrary distances on the x-axis to increase perceptibility. White circles represent medians, and horizontal bars represent the means. The shape of the violin depicts the kernel density estimate of the data.

## Discussion

The present study investigated whether higher-order associations between tasks from different memory domains — i.e., a shared temporal structure between a motor and an object sequence — facilitate memory consolidation across an overnight interval. Contrary to our hypothesis, we found no evidence that a structural overlap between motor and object sequences enhanced overnight memory consolidation in either domain. Instead, our exploratory analyses suggest that overnight gains in performance on the object task were modulated by the order of learning irrespective of the presence of higher-order associations with the motor task. Specifically, when motor sequence learning preceded object sequence learning, overnight gains in performance on the object task were impaired, reflecting pro-active interference from the earlier motor memory on object memory consolidation.

Our pre-registered analyses did not show evidence of a beneficial effect of a cross-domain shared temporal structure on memory consolidation. These results contrast with previous findings suggesting that shared structural features between tasks from different domains can promote the stabilization of memory representations [14]. Specifically, prior work has shown that when tasks share a common temporal structure, and the learning sessions are separated by sufficient time for the first memory to stabilize (as in the current study and in Brown and Robertson, 2007), the task learned second shows less susceptibility to interference from competing material [14]. This account would predict that re-exposure and rehearsal of the stabilized structure in the second learning session should further strengthen its representation [29,30]. Moreover, evidence indicates that memories actively stabilized through wakeful rehearsal are subsequently associated with greater offline gains after sleep [31,32]. Based on this framework, we expected enhanced overnight memory consolidation on both tasks when the tasks shared structural representations that were strengthened across learning episodes. The current data do not support this view. We therefore considered an alternative possibility discussed in prior research suggesting the existence of a potential cost for the first memory whereby the re-exposure to the same structure in a new context (e.g., words vs. actions) would disrupt the original memory by weakening the link between its content and the shared structure [14]. This account would predict a beneficial vs. disruptive effect of higher-order associations for the consolidation of the second task vs. the first task, respectively. We performed exploratory analyses to test this possibility (see Section 7 of the supplemental information) but did not observe such effects. Altogether, the current results therefore suggest that higher-order associations between memory domains did not influence overnight memory retention.

Interestingly, the results of our exploratory analyses indicate that while the structure of the learning experiences did not influence consolidation, their order did. Specifically, overnight gains in performance on the object task were lower when this task was learned during the second as compared to the first session. One could argue that this order effect on consolidation might be due to a general session effect, whereby performance in the second session was overall better than during the first session, which decreased the potential room for further overnight improvement. This is unlikely for the following reasons. First, negative control analyses didn’t highlight any significant session effect, i.e., average performance during session 2 was not faster than during session 1. Second, no such order effect on the consolidation metrics was observed on the motor task. Last, performance on the object task tended to deteriorate during session 2 in the associated group which – based on the hypotheses above – would have increased the potential room for improvement over the consolidation interval. Yet, we did not observe higher gains in performance on object when learnt in the second session. An alternative explanation for this pattern of results is that motor learning may have proactively interfered with the encoding of the object task on Day 1, which in turn might have compromised the subsequent object memory consolidation. While such interpretation contradicts earlier work showing that a 4h interval between learning episodes (as in the current study) is sufficient for memories from different domains to stabilize and become resistant to interference [8], it is partially in line with the results of our exploratory analyses performed on Day 1 data. Specifically, our data showed that performance on the object task was slower when learned after the motor sequence task on Day 1. However, in contrast to the effect observed on consolidation, this interfering effect tended to be more pronounced in the associated group when the two tasks shared a common temporal structure. The lack of such higher-order association effect on overnight gains in performance suggests that the association effects were only transitory. In other words, it is possible that although higher-order associations may influence initial learning, these effects did not carry over the retention interval. The dissipation of such transitory effects over a night of sleep is line with previous observations from Brown and Robertson (2007) who demonstrated that interference between memory domains typically emerges during wake consolidation but dissipates after sleep. This is also consistent with prior work suggesting that a night of sleep can restore memories previously disrupted as a result of interfering experiences the previous day [33–35]. Thus, the weaker effect of higher-order association on performance collected on Day 1 may be attenuated over the consolidation interval while the more robust proactive interference effect of the motor task on object memory consolidation might have carried overnight. It is important to note though that this is speculative and that our study was not powered to detect such effects which limits the interpretation of these results.

It remains unclear why the tasks tended to interfere despite the 4h stabilization interval between sessions on Day 1 [8]. More recent research even indicates that interaction between memory domains via high-order associations dissipates when the learning sessions are 2h apart [13].

Instead, our results suggest that memories might have remained unstable even when learning episodes are separated by 4h. Such discrepant results might be related to the nature of our tasks as they were overall more complex than in those used in previous research and may have required more time to stabilize [36]. This possibility is consistent with our observation that object learning tended to be impaired when it occurred after motor learning. However, it does not allow us to explain the lack of interference from the object to the motor task or the impact of higher-order associations on this effect. Indeed, assuming the initial memory was still unstable in the current study, previous research would predict an opposite pattern of results, i.e., transfer - rather than interference - between tasks from different domains that share a common structure [13]. Our data did not replicate these earlier observations. Notably, a recent failed replication of Mosha & Robertson (2016) reported a similar absence of transfer between tasks from different domains sharing a common structure [15]. The authors suggest that cross-domain generalization may be highly sensitive to subtle methodological variations. Differences in design in our study, such as the longer delay between learning sessions and the use of more complex visuomotor object learning rather than word lists, may have therefore contributed to the lack of transfer.

Another key discrepancy between our findings and prior research concerns the *directionality* of the interaction effects. Whereas previous work has shown that interactions between declarative and procedural memories - whether facilitative or disruptive - are typically bidirectional [8,13,14], we observed interference only in one direction. Specifically, object learning and consolidation processes were disrupted when the motor task preceded the object task. In contrast, motor learning was not affected by the preceding object learning session. This asymmetry might suggest that, in our study, the motor memory trace may be more robust than the declarative memory trace. An alternative to this *mnemonic* perspective is a *motor execution* explanation. It is possible that the task strategy developed during the motor version of the task might have interfered with the execution of the other task versions, irrespective of the memory content of the task. Specifically, with extended practice of the motor task, participants become less reliant on the visual cues to perform the series of memorized movements. One could expect that, when presented with another task version that required the use of this mapping (e.g., random or object versions of the task), participants’ performance would be greatly affected. The current data show evidence for both accounts. In line with this interference to *motor execution* hypothesis, our data show that performance on the random task (that does not carry any memory content) was slower after the motor – as compared to after the object – task session (see Section 1 of the supplements for the significant condition effect reported on the random data acquired in session 2). However - and in line with the *mnemonic* explanation – it appears that this interference remained significant during object practice in session 2 in the associated group (while it dissipated during object practice in the unassociated group, see Figure 4A-B), suggesting that the interference process went beyond simple motor execution in the associated group. The *mnemonic* alternative is also in line with the observation that object memory retention (retested overnight after the potential dissipation of the interference to execution) was affected by prior motor learning. It should be stated that these two explanations (i.e., execution vs. mnemonic) are certainly not mutually exclusive. Thus, our data suggests that motor learning prior to object learning may have resulted in both a transitory interference to execution as well as an active process of proactive interference to subsequent object learning and object memory consolidation.

In summary, our data suggest that higher-order structural associations between the procedural and the declarative memory domains did not enhance sequence learning and overnight memory consolidation. Instead, the order of the different learning episodes appeared to be a stronger determinant of the consolidation process. Our findings suggest that careful consideration should be given to the order of learning experiences when designing educational interventions or rehabilitation protocols, where optimal scheduling could minimize interference between learning episodes and maximize memory retention.

## Acknowledgment

We thank undergraduate students Krista Hamilton, Ashmita Karki, and Ryder Robins for their valuable assistance with data collection.

## Data availability

The datasets generated and analyzed, along with the analysis scripts used in the current study, will be made available in a public data repository upon publication.

## Author contributions

GA, AT & BRK designed the experiments. AT & OB conducted the experiments. GA, AT, BRK & OB contributed to acquisition and analytic tools. AT & GA analyzed the data. All authors contributed to the manuscript.

## References

[1] Cohen NJ, Squire LR. Preserved learning and retention of pattern-analyzing skill in amnesia: dissociation of knowing how and knowing that. Science 1980;210:207–10.

[2] Davachi L, DuBrow S. How the hippocampus preserves order: the role of prediction and context. Trends Cogn Sci 2015;19:92–9.

[3] Dolfen N, Reverberi S, Op de Beeck H, King BR, Albouy G. The hippocampus represents information about movements in their temporal position in a learned motor sequence. J Neurosci 2024;44:e0584242024.

[4] Gann MA, King BR, Dolfen N, Veldman MP, Chan KL, Puts NAJ, et al. Hippocampal and striatal responses during motor learning are modulated by prefrontal cortex stimulation. Neuroimage 2021;237:118158.

[5] Hsieh L-T, Gruber MJ, Jenkins LJ, Ranganath C. Hippocampal activity patterns carry information about objects in temporal context. Neuron 2014;81:1165–78.

[6] Gabrieli JD, Corkin S, Mickel SF, Growdon JH. Intact acquisition and long-term retention of mirror-tracing skill in Alzheimer’s disease and in global amnesia. Behav Neurosci 1993;107:899–910.

[7] Squire LR, Zola SM. Structure and function of declarative and nondeclarative memory systems. Proc Natl Acad Sci U S A 1996;93:13515–22.

[8] Brown RM, Robertson EM. Off-line processing: reciprocal interactions between declarative and procedural memories. J Neurosci 2007;27:10468–75.

[9] Kamal L, Celik B, Alpert G, Gonzalez S, Nguyen V, Freedberg M. Failure to reproduce the effect of procedural memory interference on wakeful consolidation of episodic memory in younger and older adults. BioRxiv 2024. 10.1101/2024.10.17.618844.

[10] Albouy G, King BR, Maquet P, Doyon J. Hippocampus and striatum: dynamics and interaction during acquisition and sleep-related motor sequence memory consolidation. Hippocampus 2013;23:985–1004.

[11] Temudo A, Dolfen N, King BR, Albouy G. The human medial temporal lobe represents memory items in their ordinal position in both declarative and motor memory domains. PLoS Biol 2025;23:e3003267.

[12] Failla A, Bracco M, Robertson EM. A common serial structure causes offline excitability changes linked to generalization between different memory types. Curr Biol 2025. 10.1016/j.cub.2025.10.063.

[13] Mosha N, Robertson EM. Unstable memories create a High-Level representation that enables learning transfer. Curr Biol 2016;26:100–5.

[14] Mutanen TP, Bracco M, Robertson EM. A common task structure links together the fate of different types of memories. Curr Biol 2020;30:2139–2145.e5.

[15] Thong S, Hendrikse J, Chong TT-J, Coxon JP. Generalisation between motor and declarative memory sequences: A conceptual replication of Mosha & Robertson (2016). BioRxiv 2025. 10.1101/2025.10.15.682521.

[16] Faul F, Erdfelder E, Lang A-G, Buchner A. G*Power 3: a flexible statistical power analysis program for the social, behavioral, and biomedical sciences. Behav Res Methods 2007;39:175–91.

[17] Beck AT, Ward CH, Mendelson M, Mock J, Erbaugh J. An inventory for measuring depression. Arch Gen Psychiatry 1961;4:561–71.

[18] Beck AT, Epstein N, Brown G, Steer RA. An inventory for measuring clinical anxiety: psychometric properties. J Consult Clin Psychol 1988;56:893–7.

[19] Buysse DJ, Reynolds CF 3rd, Monk TH, Berman SR, Kupfer DJ. The Pittsburgh Sleep Quality Index: a new instrument for psychiatric practice and research. Psychiatry Res 1989;28:193–213.

[20] Horne JA, Ostberg O. A self-assessment questionnaire to determine morningness-eveningness in human circadian rhythms. Int J Chronobiol 1976;4:97–110.

[21] Johns MW. A new method for measuring daytime sleepiness: the Epworth sleepiness scale. Sleep 1991;14:540–5.

[22] Oldfield RC. The assessment and analysis of handedness: the Edinburgh inventory. Neuropsychologia 1971;9:97–113.

[23] King BR, Dolfen N, Gann MA, Renard Z, Swinnen SP, Albouy G. Schema and motor-memory consolidation. Psychol Sci 2019;30:963–78.

[24] Brady TF, Konkle T, Alvarez GA, Oliva A. Visual long-term memory has a massive storage capacity for object details. Proc Natl Acad Sci U S A 2008;105:14325–9.

[25] Ellis BW, Johns MW, Lancaster R, Raptopoulos P, Angelopoulos N, Priest RG. The St. Mary’s Hospital sleep questionnaire: a study of reliability. Sleep 1981;4:93–7.

[26] MacLean AW, Fekken GC, Saskin P, Knowles JB. Psychometric evaluation of the Stanford Sleepiness Scale. J Sleep Res 1992;1:35–9.

[27] Dinges DF, Powell JW. Microcomputer analyses of performance on a portable, simple visual RT task during sustained operations. Behav Res Methods Instrum Comput 1985;17:652–5.

[28] Pan SC, Rickard TC. Sleep and motor learning: Is there room for consolidation? Psychol Bull 2015;141:812–34.

[29] Antony JW, Ferreira CS, Norman KA, Wimber M. Retrieval as a fast route to memory consolidation. Trends Cogn Sci 2017;21:573–6.

[30] Dudai Y. The restless engram: consolidations never end. Annu Rev Neurosci 2012;35:227–47.

[31] Himmer L, Schönauer M, Heib DPJ, Schabus M, Gais S. Rehearsal initiates systems memory consolidation, sleep makes it last. Sci Adv 2019;5:eaav1695.

[32] Brodt S, Schönauer M, Seewald A, Beck J, Erb M, Scheffler K, et al. Memory systems integration in sleep complements rapid systems consolidation in wakefulness. BioRxiv 2023. 10.1101/2023.03.09.531360.

[33] Albouy G, King BR, Schmidt C, Desseilles M, Dang-Vu TT, Balteau E, et al. Cerebral activity associated with transient sleep-facilitated reduction in motor memory vulnerability to interference. Sci Rep 2016;6. 10.1038/srep34948.

[34] Nettersheim A, Hallschmid M, Born J, Diekelmann S. The role of sleep in motor sequence consolidation: stabilization rather than enhancement. J Neurosci 2015;35:6696–702.

[35] McDevitt EA, Duggan KA, Mednick SC. REM sleep rescues learning from interference. Neurobiol Learn Mem 2015;122:51–62.

[36] Lugassy D, Herszage J, Pilo R, Brosh T, Censor N. Consolidation of complex motor skill learning: evidence for a delayed offline process. Sleep 2018;41. 10.1093/sleep/zsy123.

