## Supplementary Information for "The order, but not the structure, of cross-domain learning influences memory consolidation"

**Authors:** Ainsley Temudo, Nina Dolfen, Bradley R. King and Genevieve Albouy

**Corresponding author:** Genevieve Albouy

**This PDF file includes:**

Supporting text 1-7
Figures S1 to S3
Tables S1 to S4

### 1. Assessment of baseline performance on random SRTT

#### *Session effects*

We tested whether performance on the random SRTT was different between sessions (1 vs. 2). Performance speed on the random SRTT (Figure 3A) was faster for session 2 than for session 1 (session: $F(1, 46) = 44.74$, $p < .001$, $\eta_p^2 = .49$; session x block: $F(3, 138) = 9.78$, $p < .001$, $\eta_p^2 = .18$) while performance accuracy was greater for session 1 than for session 2 (session: $F(1, 46) = 5.69$, $p = .02$, $\eta_p^2 = .11$; session x block: $F(3, 138) = 1.15$, $p = .33$, $\eta_p^2 = .02$). This effect did not differ between groups for speed (group: $F(1, 46) = 1.53$, $p = .22$, $\eta_p^2 = .03$; session x group: $F(1, 46) = .40$, $p = .53$, $\eta_p^2 = .009$; session x block x group: $F(3, 138) = 1.55$, $p = .20$, $\eta_p^2 = .03$) or accuracy (group: $F(1, 46) = 2.61$, $p = .11$, $\eta_p^2 = .05$; session x group: $F(1, 46) = .43$, $p = .51$, $\eta_p^2 = .009$; session x block x group: $F(3, 138) = 2.98$, $p = .03$, $\eta_p^2 = .06$). Together, these results demonstrate that participants tended to be faster but less accurate in session 2 as compared to session 1. However, these effects are unlikely to affect the results presented in the main text as the order of the two conditions (motor and declarative) was counterbalanced across participants. Importantly, motor execution was in general equivalent between the associated and unassociated groups, both in terms of speed and accuracy.

#### *Condition effects*

We tested whether general motor execution, assessed with the random SRTT, differed at baseline between the associated and unassociated groups and the motor and object task conditions on Day 1. Results showed that performance speed (i.e., mean response time) and accuracy (i.e., %

correct responses) remained stable across blocks of the random SRTT (speed: $F(3, 138) = .70$, p = .56, $\eta_p^2 = .02$; accuracy: $F(3, 138) = .88$, p = .45, $\eta_p^2 = .02$) and this effect did not differ between task conditions (speed; condition: $F(1, 46) = .94$, p = .34, $\eta_p^2 = .02$; condition x block: $F(3, 138) = 1.65$, p = .18, $\eta_p^2 = .04$; accuracy; condition: $F(1, 46) = .05$, p = .82, $\eta_p^2 = .001$; condition x block: $F(3, 138) = 1.78$, p = .15, $\eta_p^2 = .04$) and importantly did not differ between groups (speed; group: $F(1, 46) = 1.53$, p = .22, $\eta_p^2 = .03$; block x group: $F(3, 138) = .17$, p = .92, $\eta_p^2 = .004$; condition x group: $F(1, 46) = .45$, p = .51, $\eta_p^2 = .01$; condition x block x group: $F(3, 138) = .16$, p = .92, $\eta_p^2 = .004$; accuracy; group: $F(1, 46) = 2.61$, p = .11, $\eta_p^2 = .05$; block x group: $F(3, 138) = .54$, p = .66, $\eta_p^2 = .01$; condition x group: $F(1, 46) = .22$, p = .64, $\eta_p^2 = .005$; condition x block x group: $F(3, 138) = .92$, p = .43, $\eta_p^2 = .02$). Together, these results confirm that baseline motor execution was statistically equivalent across groups and task conditions, both in terms of speed and accuracy.

#### *Task order effects*

We tested whether the order of conditions across sessions influenced baseline performance differently between groups. Results showed that task order did not differently influence performance between groups (speed: task order x group: $F(1, 44) = .04$, $p = .85$, $\eta p^2 = .001$; session x task order x group: $F(1, 44) = .03$, $p = .86$, $\eta p^2 = .001$; block x session x task order x group: $F(3,132) = .51$, $p = .67$, $\eta p^2 = .01$; accuracy: task order x group: $F(1, 44) = .12$, $p = .73$, $\eta p^2 = .003$; session x task order x group: $F(1, 44) = .21$, $p = .65$, $\eta p^2 = .005$; block x session x task order x group: $F(3, 132) = 1.12$, $p = .34$, $\eta p^2 = .03$ ). Even though the order of the different conditions did not affect performance differently between groups, we ran exploratory analyses to further examine condition effects within each sessions.

A block x condition (motor vs object) x group RM ANOVA conducted on the random SRTT of **session 1,** showed that performance speed improved across blocks (block: $F(3, 132) = 13.34$, $p < .001$, $\eta_p^2 = .23$) and was faster when it was followed by the object task compared to when it was followed by the motor task (condition: $F(1, 44) = 4.48$, $p = .04$, $\eta_p^2 = .09$; condition x block: $F(3, 132) = .02$, $p < 1$, $\eta_p^2 = .00$) but did not differ between groups (group: $F(1, 44) = 4.48$, $p = .04$, $\eta_p^2 = .09$; block x group: $F(3, 132) = 1.50$, $p = .22$, $\eta_p^2 = .03$; condition x group: $F(1, 44) = .27$, $p = .61$, $\eta_p^2 = .006$; group x block x condition: $F(3, 132) = .35$, $p = .79$, $\eta_p^2 = .008$). Performance accuracy remained stable across blocks (block: $F(3, 132) = 1.33$, $p = .27$, $\eta_p^2 = .03$) and did not differ between conditions (condition: $F(1, 44) = .27$, $p = .61$, $\eta_p^2 = .006$; condition x block: $F(3, 132) = .39$, $p = .76$, $\eta_p^2 = .009$ ) or groups (group: $F(1, 44) = 1.96$, $p = .17$, $\eta_p^2 = .04$; block x group: $F(3, 132) = 1.71$, $p = .17$, $\eta_p^2 = .04$; condition x group: $F(1, 44) = .35$, $p = .56$, $\eta_p^2 = .008$; group x block x condition: $F(3, 132) = .94$, $p = .42$, $\eta_p^2 = .02$).

The block x condition (motor vs object) x group RM ANOVA conducted on the random SRTT of **session 2** showed that performance remained stable across blocks (block; speed: $F(3, 132) = 2.66$, $p = .06$, $\eta_p^2 = .06$; accuracy: $F(3, 132) = 1.33$, $p = .27$, $\eta_p^2 = .03$), but was slower when it preceded the object task compared to when it preceded the motor task (speed; condition: $F(1, 44) = 12.97$, $p < .001$, $\eta_p^2 = .23$; condition x block: $F(3, 132) = 2.72$, $p = .05$, $\eta_p^2 = .06$). In other words, these data suggest that learning the motor task in session 1 resulted in slower performance on the random task during session 2. However, this effect did not differ between groups (speed; group:

$F(1, 44) = 1.08$, p = .31, $\eta_p^2 = .09$; block x group: $F(3, 132) = .68$, p = .57, $\eta_p^2 = .02$; condition x group: $F(1, 44) = .05$, p = .83, $\eta_p^2 = .001$; group x block x condition: $F\ (3, 132) = .16$, p = .93, $\eta_p^2 = .004$). As for accuracy, performance was similar whether it was followed by the motor or object task (condition: $F(1, 44) = .32$, p = .57, $\eta_p^2 = .007$; condition x block: $F(3, 132) = 1.48$, p = .22, $\eta_p^2 = .03$) and was also similar between groups (group: $F(1, 44) = 2.34$, p = .13, $\eta_p^2 = .05$; block x group: $F(3, 132) = 1.31$, p = .27, $\eta_p^2 = .03$; condition x group: $F(1, 44) = .01$, p = .91, $\eta_p^2 = .00$; group x block x condition: $F(3, 132) = .60$, p = .62, $\eta_p^2 = .01$). Overall, these results support the *motor execution* hypothesis discussed in the main text whereby performance speed on the random task is impaired when performed after the motor task as compared to after the object task session.

### 2. Sequence-specific learning

To assess sequence-specific learning, performance was compared between the motor and object sequence conditions and the random condition at retest. For speed, an effect of condition was observed (condition: $F(1, 46) = 217.15$, $p < .001$, $\eta_p^2 = .83$; condition x block: $F(6, 276) = 3.01$, $p = .007$, $\eta_p^2 = .06$) such that speed was faster for the motor task compared to the object task (p < .001) and for both sequence tasks (motor and object) compared to the random condition (motor vs. random: p < .001; object vs. random: p < .001). This effect was not different between the associated and unassociated groups (group: $F(1, 46) = .17$, $p = .68$, $\eta_p^2 = .004$; condition x group: $F(1, 46) = 1.71$, $p = .19$, $\eta_p^2 = .04$; condition x block x group: $F(6, 276) = .72$, $p = .64$, $\eta_p^2 = .02$).

Regarding accuracy, a sequence-specific effect was also observed (condition: $F(1, 46) = 35.26$, $p < .001$, $\eta_p^2 = .43$; condition x block: $F(6, 276) = .84$, $p = .54$, $\eta_p^2 = .02$) where performance was more accurate for the motor task compared to the object task ($p < .001$) and for both sequence tasks compared to the random condition (motor vs. random: $p < .001$; object vs. random: $p = .001$). Again, this effect did not differ between the two groups (group: $F(1, 46) = .83$, $p = .37$, $\eta_p^2 = .02$; condition x group: $F(1, 46) = 1.25$, $p = .29$, $\eta_p^2 = .03$; condition x block x group: $F(6, 276) = .47$, $p = .83$, $\eta_p^2 = .01$). Altogether these results suggest that performance on the sequence tasks at retest reflect sequence-specific learning rather than general enhancement in motor execution and did not differ between associated and unassociated groups.

### 3. Effect of higher-order associations on offline changes in performance accuracy

Our main confirmatory analysis aimed to assess whether performance gains differed depending on the presence of a common learning structure between a motor and object sequence. As pre-registered, performance accuracy was compared from the end of training on Day 1 to retest on Day 2 between the two groups (associated and unassociated) irrespective of task condition (Figure S1B). Results showed a significant effect of session, whereby accuracy improved from the end of training to retest ($F(1, 46) = 10.58$, $p = .002$, $\eta p^2 = .19$). In contrast to our expectations, this session effect did not differ between the 2 groups (session x group: $F(1, 46) = .68$, $p = .42$, $\eta p^2 = .01$; group: $F(1, 46) = .12$, $p = .74$, $\eta p^2 = .003$), indicating that the presence of higher-order associations between sequences from the two different domains did not enhance offline memory consolidation as assessed with performance accuracy.

### 4. Negative control analyses on performance accuracy

To ensure that any observed effects in offline changes in performance accuracy were not influenced by differences in performance during initial training, we tested for potential group effects during initial task practice on Day 1.

Results of the group x session (session 1 vs. session 2, irrespective of task condition) x block repeated-measures ANOVA (Figure S1A) showed that accuracy remained stable across blocks (block; training: $F(19, 874) = .79$, $p = .72$, $\eta p^2 = .02$; test; $F(3, 138) = 1.65$, $p = .18$, $\eta p^2 = .04$). As expected, performance accuracy did not differ between the associated and unassociated groups (training; group: $F(1, 46) = 2.09$, $p = .16$, $\eta p^2 = .04$; block x group: $F(19, 874) = .51$, $p = .96$, $\eta p^2 = .01$; test; group: $F(1, 46) = .003$, $p = .96$, $\eta p^2 = .00$; block x group: $F(3, 138) = .14$, $p = .94$, $\eta p^2 = .003$) but did differ between sessions during training (not at test). Specifically, accuracy was higher during session 1 compared to session 2 (training; session: $F(1, 46) = 5.65$, $p = .02$, $\eta p^2 = .11$; session x block: $F(19, 874) = .75$, $p = .77$, $\eta p^2 = .02$; test; session: $F(1, 46) = .03$, $p = .87$, $\eta p^2 = .001$; session x block: $F(3, 138) = 2.23$, $p = .09$, $\eta p^2 = .05$). No significant interactions were observed between these factors (training; session x group: $F(1, 46) = .003$, $p = .96$, $\eta p^2 = .00$; session x block x group: $F(19, 874) = .68$, $p = .84$, $\eta p^2 = .01$; test; session x group: $F(1, 46) = 3.25$, $p = .08$, $\eta p^2 = .07$; session x block x group: $F(3, 138) = .60$, $p = .62$, $\eta p^2 = .01$).

The group x condition (motor vs. object, irrespective of session) x block repeated-measures ANOVA showed a significant condition effect whereby accuracy was higher for the motor task compared to the object task during training (training; condition: $F(1, 46) = 15.17$, $p < .001$, $\eta p^2 =$

.25; condition by block: $F(19, 874) = 2.22$, $p = .002$, $\eta p^2 = .05$; test; condition: $F(1, 46) = 3.53$, $p = .07$, $\eta p^2 = .07$; condition x block: $F(3, 138) = .64$, $p = .59$, $\eta p^2 = .01$). However, this effect did not interact with the factor group (training; condition x group: $F(1, 46) = .001$, $p = .98$, $\eta p^2 = .00$; condition x block x group: $F(19, 874) = 1.25$ $p = .21$, $\eta p^2 = .03$; test; condition x group: $F(1, 46) = .09$, $p = .76$, $\eta p^2 = .002$; condition x block x group: $F(3, 138) = .25$ $p = .86$, $\eta p^2 = .005$)

We then tested whether the order of motor and object task conditions across sessions influenced accuracy differently between the associated and unassociated groups. To do so, a group x session x order (motor first vs. object first) x block repeated-measures ANOVA was performed. Results showed that task order did not differently influence performance accuracy between groups (training; order x group: $F(1, 44) = .31$, $p = .58$, $\eta p^2 = .007$; order x group x session: $F(1, 44) = .001$, $p = .98$, $\eta p^2 = .00$; order x group x session x block: $F(19, 836) = 1.23$, $p = .22$, $\eta p^2 = .03$; test; order x group: $F(1, 44) = .02$, $p = .90$, $\eta p^2 = .00$; order x group x session: $F(1, 44) = .10$, $p = .76$, $\eta p^2 = .002$; order x group x session x block: $F(3, 132) = .25$, $p = .86$, $\eta p^2 = .006$ ).

Overall, these negative control analyses indicate that accuracy did not differ between groups during initial learning but that participants were more accurate on the motor compared to object task and more accurate in the first session compared to the second. Even though the order of these different tasks did not affect accuracy differently between groups, as in the analyses for speed, we ran exploratory analyses to further examine these effects within each session separately.

### 5. Exploratory analyses on performance accuracy

#### *Within-session analyses (Day 1)*

The block x condition x group RM ANOVA conducted on ***session 1*** (Figure S2) data confirmed a significant condition effect, whereby accuracy for the motor task was higher when compared to the object task during training (training; condition: $F(1, 44) = 14.07$, $p < .001$, $\eta_p^2 = .24$; block x condition: $F(19, 836) = 1.99$, $p = .007$, $\eta_p^2 = .04$; test; condition: $F(1, 44) = .29$, $p = .59$, $\eta_p^2 = .007$; block x condition: $F(3, 132) = .69$, $p = .56$, $\eta_p^2 = .02$). However, this effect did not differ between the associated and unassociated groups (training; group: $F(1, 44) = 2.47$, $p = .12$, $\eta_p^2 = .05$; condition x group: $F(1, 44) = .42$, $p = .52$, $\eta_p^2 = .01$; block x group: $F(19, 836) = .65$, $p = .87$, $\eta_p^2 = .02$; block x group x condition: $F(19, 836) = 1.02$, $p = .44$, $\eta_p^2 = .02$; test; group: $F(1, 44) = .70$, $p = .41$, $\eta_p^2 = .02$; condition x group: $F(1, 44) = .06$, $p = .81$, $\eta_p^2 = .001$; block x group: $F(3, 132) = .34$, $p = .79$, $\eta_p^2 = .008$; block x group x condition: $F(3, 132) = .48$, $p = .70$, $\eta_p^2 = .01$).

In contrast, no condition effect was observed in **session 2** (Figure S2; training; condition: $F(1, 44) = 1.39$, $p = .24$, $\eta_p^2 = .03$; block x condition: $F(19, 836) = 1.42$, $p = .11$, $\eta_p^2 = .03$; test: condition; $F(1, 44) = 1.58$, $p = .22$, $\eta_p^2 = .04$; block x condition: $F(3, 132) = .19$, $p = .91$, $\eta_p^2 = .004$) and no group effects or interactions with group were observed during training (group: $F(1, 44) = 1.11$, $p = .30$, $\eta_p^2 = .03$; condition x group: $F(1, 44) = .15$, $p = .70$, $\eta_p^2 = .003$; block x group: $F(19, 836) = .58$, $p = .92$, $\eta_p^2 = .01$; block x group x condition: $F(19, 836) = 1.13$, $p = .31$, $\eta_p^2 = .03$) or at test ( group: $F(1, 44) = .72$, $p = .40$, $\eta_p^2 = .02$; condition x group: $F(1, 44) = .001$, $p = .97$, $\eta_p^2 = .00$; block x group: $F(3, 132) = .42$, $p = .74$, $\eta_p^2 = .009$; block x group x condition: $F(3, 132) = 1.49$, $p = .22$, $\eta_p^2 = .03$). Overall, these findings suggest that higher-order associations did not influence performance accuracy during the acquisition of the tasks learned in either session.

### 6. Generation

Analyses of the generation tasks demonstrated that participants successfully reproduced the motor and object sequences with high accuracy. Specifically, the mean percentage of correct ordinal positions, i.e., the percentage of items (movements or objects) generated in the correct temporal position in the sequence and the percentage of correct transitions generated was greater than 89% across tasks (% correct ordinal; motor: 95% (12.1), obj: 89% (26.2); % correct transitions; motor: 94% (13.3), object: 91% (22.8)). A repeated-measures ANOVA revealed no significant differences between object and motor sequence generation scores in terms of both correct ordinal positions (condition: $F(1, 46) = 3.15$, $p = .08$, $\eta_p^2 = .06$) and correct transitions (condition: $F(1, 46) = 1.62$, $p = .21$, $\eta_p^2 = .03$). Importantly, no significant differences in generation scores were observed between groups for correct ordinal positions (group: $F(1, 46) = 1.45$, $p = .24$, $\eta_p^2 = .03$; condition x group: $F(1, 46) = .69$, $p = .41$, $\eta_p^2 = .02$), nor correct transitions (group: $F(1, 46) = .57$, $p = .46$, $\eta_p^2 = .01$; condition x group: $F(1, 46) = .03$, $p = .87$, $\eta_p^2 = .001$). These results indicate that participants from both groups developed explicit knowledge of the series of objects and movements.

### 7. Beneficial vs. disruptive effect

We tested whether higher-order associations would produce a disruptive effect on the first learning task and a beneficial effect on the second task, as suggested in previous research [14]. To evaluate this, we conducted independent-samples t-tests comparing the associated and unassociated groups on: (1) gains from session 1 irrespective of task condition—where a disruptive effect would predict lower gains in the associated group compared to the unassociated group—and (2) gains from session 2 irrespective of task condition—where a beneficial effect would predict higher gains in the associated group compared to the unassociated group. No significant group effects were observed for either session 1 gains ($t(46) = -0.17$, $p = .43$) or session 2 gains ($t(46) = -0.27$, $p = .39$). These results indicate that introducing higher-order associations did not produce a trade-off between tasks where a strengthening of the second memory came at the expense of the first task.

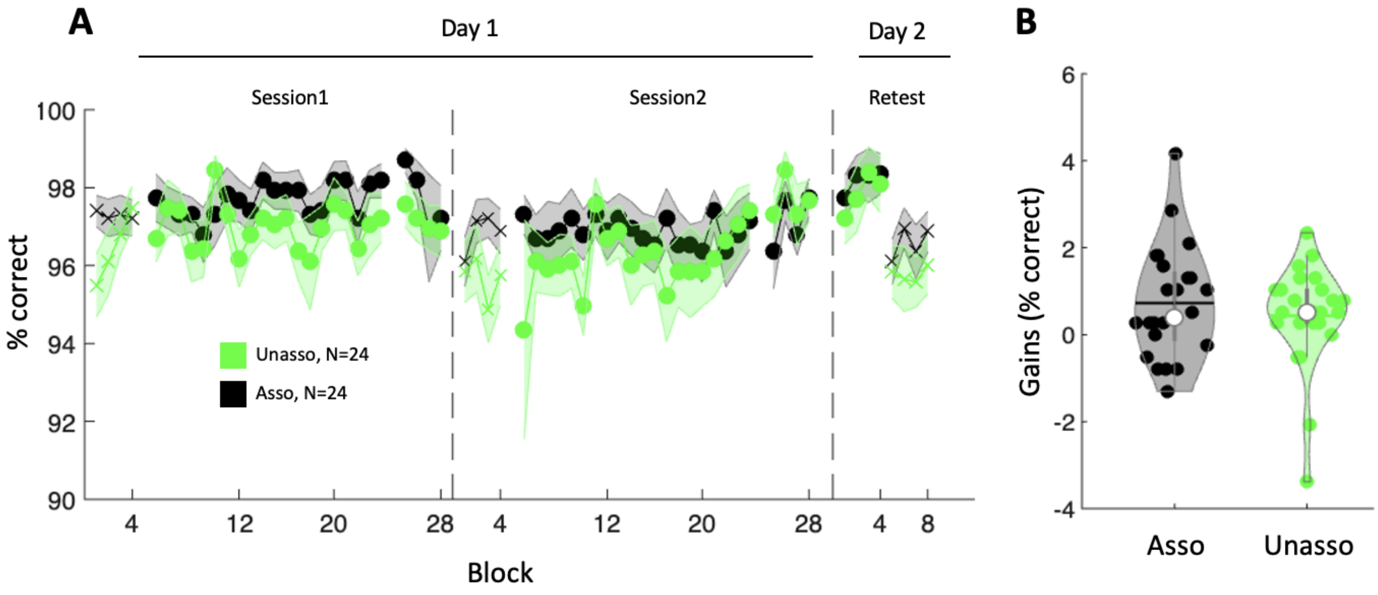


*Figure S1*. *Performance accuracy results* **A**. Accuracy (% correct responses) across blocks and sessions collapsed across motor and object task conditions for the unassociated ("Unasso", green, N = 24) and associated ("Asso", black, N = 24) groups. Crosses and circles indicate blocks of the random and sequence conditions, respectively. Shaded error bars=SEM. **B**. Gains in performance accuracy calculated as the average accuracy across the four test blocks on Day 1 minus the average across the four retest blocks on Day 2, for each group. Gains were not significantly different between the associated and unassociated group indicating that higher-order associations between sequences across different memory domains did not enhance offline memory consolidation. Colored circles represent individual data, jittered in arbitrary distances on the x-axis to increase perceptibility. White circles represent medians, and horizontal bars represent means. The shape of the violin depicts the kernel density estimate of the data.

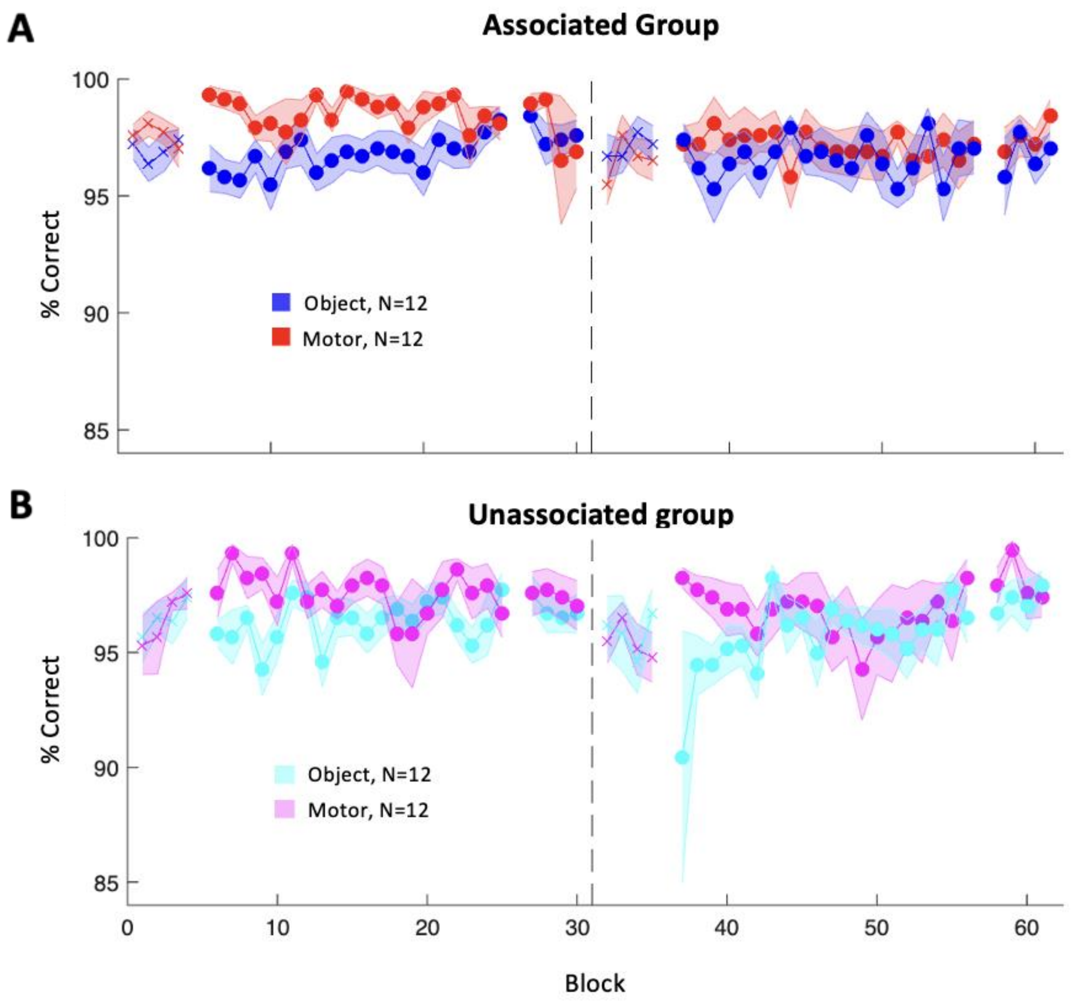


*Figure S2*. *Results of the exploratory analyses on performance accuracy* **A**. Accuracy (% of correct responses) across blocks in session 1 and 2 for the object condition (dark blue, N = 12) and motor condition (red, N = 12) in the associated group. **B**. Response time across blocks in session 1 and 2 for the object (cyan, N = 12) and motor condition (pink, N = 12) in the unassociated group. Crosses indicate blocks of the random condition; circles indicate blocks of the sequence condition. Shaded error bars=SEM.

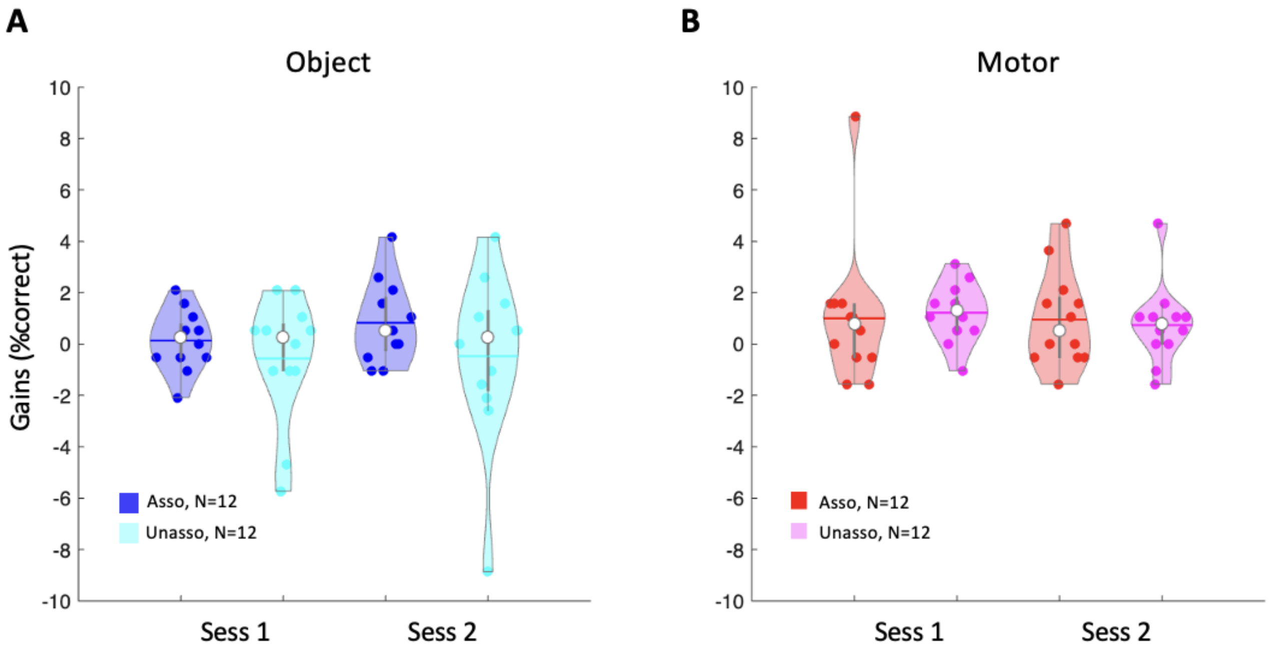


*Figure S3*. *Gains in Accuracy*. **A**. Gains on the object task in each group depending on whether the object task was learned during session 1 or session 2. **B.** Gains on the motor task in each group depending on whether the motor task was learned during session 1 or session 2. In both panels, gains were calculated as the average response time across the four test blocks on Day 1 minus the average across the four retest blocks on Day 2. Gains in performance accuracy were not significantly different between groups or session for either task condition. Colored circles represent individual data, jittered in arbitrary distances on the x-axis to increase perceptibility. White circles represent medians, and horizontal bars represent the means. The shape of the violin depicts the kernel density estimate of the data.

| N (48) | Associated (24) | Unassociated (24) | Unpaired t-test/Chi squared |
|---|---|---|---|
| Age (yrs) | 23 ± 4.1 | 21.1 ± 3.1 | t(46) = 1.8; p = 0.08, d = 0.5 |
| Sex | 14(F) 8(M) 2(NB) | 16(F) 8(M) | χ2 (2, N = 48) = 2.1, p = 0.3 |
| *a* Edinburgh Handedness (Oldfield, 1971) | 68.5 ± 45 | 70.8 ± 31.3 | t(46) = -0.2; p = 0.8, d = -0.1 |
| *b* Epworth Sleepiness Scale (Hoddes et al., 1972) | 5.29 ± 3.5 | 5.29 ± 2.7 | t(46) = 0; p < 1, d = 0 |
| *c* Beck Depression Scale (Beck et al., 1996) | 4.5 ± 3 | 4.25 ± 4.3 | t(46) = 0.2; p = 0.8, d = 0.1 |
| *d* Beck Anxiety Scale (Beck et al., 1988) | 4.5 ± 3.5 | 5.3 ± 4.6 | t(46) = -0.7; p = 0.5, d = -0.2 |
| *e* PSQI (Buysse et al., 1989) | 3.6 ± 1.7 | 4.2 ± 2.2 | t(46) = -1; p = 0.3, d = -0.3 |
| *f* Chronoscore (CRQ) (Horne & Östberg, 1976) | 51.3 ± 6.5 | 50 ± 7.5 | t(46) = 0.7; p = 0.5, d = 0.2 |
| ***g* St. Mary's questionnaire** | | | |
| Duration (Night preceding day 1) | 7.5 ± 0.8 | 7.5 ± 0.6 | t(46) = -0.3; p = 0.7, d = -0.1 |
| Duration (Night preceding day 2) | 7.8 ± 1.1 | 7.6 ± 1.4 | t(46) = 0.7; p = 0.5, d = 0.2 |
| Quality (Night preceding day 1) | 3.9 ± 0.9 | 3.9 ± 1.1 | t(46) = 0.6; p = 0.6, d = 0.2 |
| Quality (Night preceding day 2) | 3.9 ± 0.7 | 3.5 ± 0.9 | t(46) = 1.8; p = 0.08, d = 0.5 |
| ***h* Sleep diary/Actigraph** | | | |
| Duration (Night preceding day 2) | 8.3 ± 1.1 | 8.1 ± 0.9 | t(46) = 0.5; p = 0.6, d = 0.1 |

*Table S1. Participant demographics and sleep characteristics*. ***Notes.*** Values are means and standard deviations. Duration unit is in hours. Statistical analyses showed no significant group differences for all variables. PSQI = Pittsburgh Sleep Quality Index. CRQ = Circadian Rhythm Questionnaire. [a] -100 = completely left-handed, 100 = completely right-handed; [b] 0 = normal, 24 = severe sleepiness [cut off < 10]; [c] 0 = no depression 63 = severe depression [cut off < 17]; [d] 0 = minimal anxiety, 63 = severe anxiety [cut off < 17]; [e] 0 = good sleep quality, 21 = poor sleep quality [cut off < 8]; [f] 16 = extreme evening type, 86 = extreme morning type [cut off 30 < n < 70]; [g] 1 = very unsatisfied, 5 = completely satisfied; [h] Determined in combination between sleep diary and wrist actigraphy recordings on night 4 (between the two experimental sessions).

| Variable | Associated (24) | Unassociated (24) | Factors | Stats |
|---|---|---|---|---|
| **Sleep duration (Sleep diary)** | | | | |
| Night 1 | 8.3 ± 1.1 | 8.4 ± 0.8 | | |
| Night 2 | 8.5 ± 1.6 | 8.5 ± 1.2 | Night x Group | F(3,138) = 0.2; p = 0.9, ηp2 = 0.01 |
| Night 3 | 8.2 ± 1.0 | 8.0 ± 0.7 | Night | F(3,138) = 2.2; p = 0.1, ηp2 = 0.05 |
| Night 4 | 8.5 ± 1.3 | 8.4 ± 0.8 | Group | F(1,46) = 0.05; p = 0.8, ηp2 = 0.001 |
| ***a*** **Psychomotor Vigilance Task** | | | | |
| Session 1 | 0.3 ± 0.1 | 0.3 ± 0.04 | Session x Group | F(2,90) = 0.8; p = 0.5, ηp2 = 0.02 |
| Session 2 | 0.3 ± 0.1 | 0.3 ± 0.1 | Session | F(2,90) = 1.5; p = 0.2, ηp2 = 0.03 |
| Session 3 | 0.3 ± 0.1 | 0.3 ± 0.1 | Group | F(1,45) = 0.5; p = 0.5, ηp2 = 0.01 |
| ***b*** **Stanford sleepiness score** | | | | |
| Session 1 | 2.3 ± 1.0 | 2.2 ± 0.7 | Session x Group | F(2,92) = 2.1; p = 0.1, ηp2 = 0.04 |
| Session 2 | 2.2 ± 1.1 | 1.6 ± 0.7 | Session | F(2,92) = 4.3; p = 0.02, ηp2 = 0.1 |
| Session 3 | 1.8 ± 1.1 | 1.8 ± 0.8 | Group | F(1,46) = 1.5; p = 0.2, ηp2 = 0.03 |

*Table S2. Sleep durations (according to sleep log and vigilance/sleepiness scores).* Values are means and standard deviations. Repeated-measures ANOVAs showed no significant group differences. Night 1-3 refers to sleep durations prior to the first experimental session, Night 4 refers to the sleep duration between session 2 and 3. Session 1 and 2 were performed on Day 1, 4h apart. Session 3 was performed approximately 24 h later. [a] Reaction times are in seconds (objective vigilance measure), [b] Subjective vigilance score, 1 = more alert, 7 = less alert.

| **Factors** | **Motor** | | | **Object** | | |
|---|---|---|---|---|---|---|
| **Train** | F | P | $\eta_p^2$ | F | P | $\eta_p^2$ |
| block | 25.12 | **<.001** | 0.36 | 29.62 | **<.001** | 0.40 |
| block x group | 0.36 | < 1 | 0.01 | 1.58 | **0.05** | 0.04 |
| block x seq | 1.34 | 0.15 | 0.03 | 0.72 | 0.80 | 0.02 |
| block x group x seq | 0.52 | 0.95 | 0.01 | 1.27 | 0.20 | 0.03 |
| group | 0.87 | 0.36 | 0.02 | 0.10 | 0.75 | 0.00 |
| seq | 2.29 | 0.14 | 0.05 | 0.10 | 0.75 | 0.00 |
| group x seq | 0.19 | 0.66 | 0.00 | 0.11 | 0.75 | 0.00 |
| **Test** | | | | | | |
| block | 3.67 | **0.01** | 0.08 | 0.36 | 0.78 | 0.01 |
| block x group | 0.98 | 0.40 | 0.02 | 0.24 | 0.87 | 0.01 |
| block x seq | 0.55 | 0.65 | 0.01 | 0.60 | 0.61 | 0.01 |
| block x group x seq | 0.43 | 0.74 | 0.01 | 0.27 | 0.85 | 0.01 |
| group | 0.01 | 0.92 | 0.00 | 0.32 | 0.57 | 0.01 |
| seq | 0.59 | 0.45 | 0.01 | 0.15 | 0.70 | 0.00 |
| group x seq | 0.82 | 0.37 | 0.02 | 0.09 | 0.77 | 0.00 |

*Table S3. Sequence type statistics for reaction time***.** Results of repeated-measures ANOVA with within subject factor block (1-4) and between subject factors group (associated, unassociated), and sequence type (A or B)**.** Analysis performed separately for training and test phases and for object and motor conditions. Results show no significant difference between sequence A and B in either task condition or group. Bold values = significant values.

| Factors | Motor | | | Object | | |
|---|---|---|---|---|---|---|
| **Train** | F | P | $\eta_p^2$ | F | P | $\eta_p^2$ |
| block | 1.17 | 0.28 | 0.03 | 1.69 | **0.03** | 0.04 |
| block x group | 0.81 | 0.70 | 0.02 | 0.91 | 0.57 | 0.02 |
| block x seq | 1.10 | 0.34 | 0.03 | 1.20 | 0.25 | 0.03 |
| block x group x seq | 0.89 | 0.59 | 0.02 | 1.59 | **0.05** | 0.04 |
| group | 1.18 | 0.28 | 0.03 | 1.98 | 0.17 | 0.04 |
| seq | 1.04 | 0.31 | 0.02 | 2.24 | 0.14 | 0.05 |
| group x seq | 0.14 | 0.71 | 0.00 | 0.57 | 0.46 | 0.01 |
| **Test** | | | | | | |
| block | 1.61 | 0.19 | 0.04 | 0.38 | 0.77 | 0.01 |
| block x group | 0.27 | 0.85 | 0.01 | 0.09 | 0.97 | 0.00 |
| block x seq | 0.46 | 0.71 | 0.01 | 0.69 | 0.56 | 0.02 |
| block x group x seq | 3.74 | **0.01** | 0.08 | 0.86 | 0.47 | 0.02 |
| group | 0.01 | 0.93 | 0.00 | 0.05 | 0.83 | 0.00 |
| seq | 0.82 | 0.37 | 0.02 | 0.08 | 0.77 | 0.00 |
| group x seq | 2.22 | 0.14 | 0.05 | 0.13 | 0.72 | 0.00 |

*Table S4. Sequence type statistics for accuracy.* Results of repeated-measures Anova with within subject factor block (1-4) and between subject factors group (associated, unassociated), and sequence type (A or B)**.** Analysis performed separately for training and test phases and for object and motor conditions. Results show that generally (except for in a few blocks) there was no significant difference between sequence A and B in either task condition or group. Bold values = significant values.
